# Sustaining Control and Agency Under Threat: Computational Pathways to Persistence and Escape

**DOI:** 10.64898/2026.04.08.717273

**Authors:** Nadja R. Ging-Jehli, Russell K. Childers

## Abstract

**Significance Statement:** Adaptive behavior depends on knowing when to persist and when to let go; even when letting go appears as avoidance. While classical accounts of avoidance emphasize reward-effort trade-offs, we show that these decisions are critically guided by meta-control and inferences about outcome controllability and agency. Using a novel paradigm, we dissociate drivers of avoidance and demonstrate that threat does not uniformly promote disengagement. When outcome control is preserved, threat instead increases persistence, particularly following experiences that build agency in failure-safe contexts. We formalize these dynamics in the Meta-Arbitration of Control and Agency Q-learning (MACA-Q) model, which captures how experience-dependent beliefs about agency guide learning and choice across contexts. Our results show that similar avoidance behaviors can arise from distinct computational pathways. This shifts the focus from global avoidance biases to the dynamic regulation of agency as a core principle of adaptive behavior, with implications for neuroscience, psychiatry, and adaptive artificial intelligence.

Adaptive behavior requires deciding when to persist and when to disengage under uncertainty and partial outcome control. Avoidance has often been studied as a response to threat or cost, yet existing paradigms cannot disentangle whether disengagement reflects threat sensitivity, expected failure, or reduced perceived control.

We introduce a persistence-escape paradigm that independently manipulates incentive structures, effort demands, and outcome controllability. In a large online sample (N = 457), we show that avoidance is context-dependent rather than a stable, global trait. When outcome control was preserved under threat, the typical avoidance response reversed, promoting persistence rather than withdrawal. At the individual level, high-performing individuals were not uniformly more persistent, but more selective, disengaging when control was low. Moreover, higher anxiety symptoms were linked to cost-dominant evaluation and reduced use of accumulated competence. Conversely, higher depressive symptoms were linked to diminished sensitivity to effort and higher expected failure.

To explain these behavioral patterns, we developed the Meta-Arbitration of Control and Agency Q-learning (MACA-Q) model, which embeds value learning and affective evaluation within a meta-control architecture. Critically, we formalize *agency* as a dynamically inferred learning gate, distinct from self-efficacy, that determines whether outcomes are treated as informative based on controllability and feedback reliability. The model explains context-specific avoidance and reveals that similar behaviors can arise from distinct computational pathways. It further shows how experience in failure-safe contexts guides subsequent behavior in adverse contexts.

Our findings show that avoidance is guided by the dynamic regulation of engagement based on inferred controllability and competence. By combining a novel paradigm with a computational model, we provide a formal account of agency and a unifying framework in which meta-control regulates adaptive and maladaptive engagement across contexts, with implications for neuroscience, psychiatry, and adaptive artificial intelligence.

## Introduction

> *“All we have to decide is what to do with the time that is given us.”*

> *J.R.R. Tolkien, The Fellowship of the Ring*^1^

In an increasingly unpredictable world, we are confronted with situations where demands are constantly shifting, and outcomes are only partially controllable. How do we decide when to persist and when to let go? When is disengagement an adaptive response to uncertainty, and when does it become a rigid pattern of avoidance that undermines goal-directed behavior? Answering these questions have become pressing in the context of political and economic instability, and unexpected societal changes. Under these conditions, we are repeatedly required to effectively decide how to act with the time, control, and resources available to us.

Decisions between persistence and escape require arbitration over the relative benefits and costs of each alternative.^2–5^ We use the term *escape* to describe the decision to disengage from an imminent or ongoing task, without presupposing its motivational or affective drivers. Existing research often conflates the biological and psychological mechanisms of persistence and escape with those of approach and avoidance.^6–8^ However, some studies indicate that these behaviors recruit fundamentally different modes of goal-directed control.^9,10^ While escape can stem from avoidance of anticipated negative outcomes,^11^ it may also represent an adaptive response to low outcome controllability or idiosyncratic cost-benefit trade-offs.^5,12–14^ Similarly, while persistence can reflect the simple pursuit of high-value rewards, it often requires active mechanisms to overcome accumulated effort costs, setbacks, and to maintain beliefs of outcome controllability. Distinguishing these actions from their traditional functional interpretations is therefore critical for understanding when escapism reflects maladaptive avoidance versus adaptive reallocation of effort.^8,15,16^

There is extensive interest across neuroscience,^2,17^ psychology,^3,18^ psychiatry,^13,19,20^ and economics^21–23^ in the situational and psychological factors underlying avoidance behavior (including escapism) given its central role in learning,^5,17^ decision-making,^18,24^ and mental health.^13,25^ Across disciplines and species, avoidance has been shown to increase under threat,^8,26^ but it is increasingly acknowledged to not only represent reflexive responses.^27,28^ Instead, it can also emerge from cost-benefit evaluations that integrate anticipated rewards, effort demands, uncertainty and expected outcomes.^14,27,29–32^ To date, most paradigms remain unsuited to capture how individuals arbitrate between persistence and escape in more nuanced settings that closer represent characteristics of real-world settings.^8,15,33–35^ Instead, approach-avoidance behavior is often studied in isolation,^15,19^ collapsing failures of persistence with strategic escape, and conflating effort costs with feasibility.^33,36^ In particular, paradigms rarely manipulate controllability over uncertain outcomes or account for the economic, psychological, and self-evaluative costs of escapism.^8,15^ This limits insights into underlying dynamic trade-offs that determine when escape reflects flexible, adaptive avoidance.

Avoidance is a prominent feature of anxiety and depression,^13,29,37^ and more broadly can characterize how individuals adapt to uncertainty in everyday life.^38–40^ Converging evidence suggests that avoidance is not a unitary construct but rather context-dependent,^13,28^ arising from distinct underlying mechanisms.^8,41^ In particular, avoidance can reflect both reflexive responses to threat (commonly linked to anxiety) and learned biases regarding agency (commonly linked to depression). For instance, anxiety is often associated with increased intolerance of uncertainty and potential costs,^8,42,43^ leading to generalized avoidance and inflexibility to adapt to contextual change.^20,44^ Conversely, depression is closely associated with *learned helplessness*,^9,12^ where experiences of low controllability reduce persistence through altered inference about agency and competency.^45^ Existing paradigms often report global avoidance biases associated with anxiety and depressive symptoms, but fail to dissociate whether withdrawal reflects threat sensitivity, reduced perceived agency due to loss of control, or feasibility- and competence-related expectations of failure.^8,33^ As a result, distinct underlying sources of similar avoidance behavior remain indistinguishable within a given context, yet give rise to diverging behavioral trajectories when task demands change.^8,15^

To understand the mechanisms underlying avoidance, we need a paradigm that separates persistence from escape, dissociates effort costs from feasibility, and systematically varies both relative reward and outcome controllability. Such a paradigm also allows us to integrate constructs (e.g., effort-reward trade-offs, cognitive control, reinforcement learning) that are typically studied in isolation across psychology, neuroscience, and decision science.^15,35,46^ By relating these constructs within a unified paradigm, we can better understand how their interplay guides behavior. This enables us to dissociate distinct pathways into avoidance and to distinguish context-sensitive escape from rigid, generalized avoidance.

For this study, we developed a novel task structure that formalizes avoidance as an explicit choice between persistence and escape under varying task contexts (Fig 1A-D). Rather than inferring avoidance indirectly from performance, our paradigm renders escape an explicit alternative whose advantage depends on three contextual factors (Fig 1C): the incentive regime (safe versus threat contexts), the trial-wise value difference between escape and persistence, and outcome controllability (the degree to which outcomes depend on action). This structure dissociates trial-level sensitivity to costs and benefits of actions from context-dependent shifts in decision policy.

**Fig. 1.**
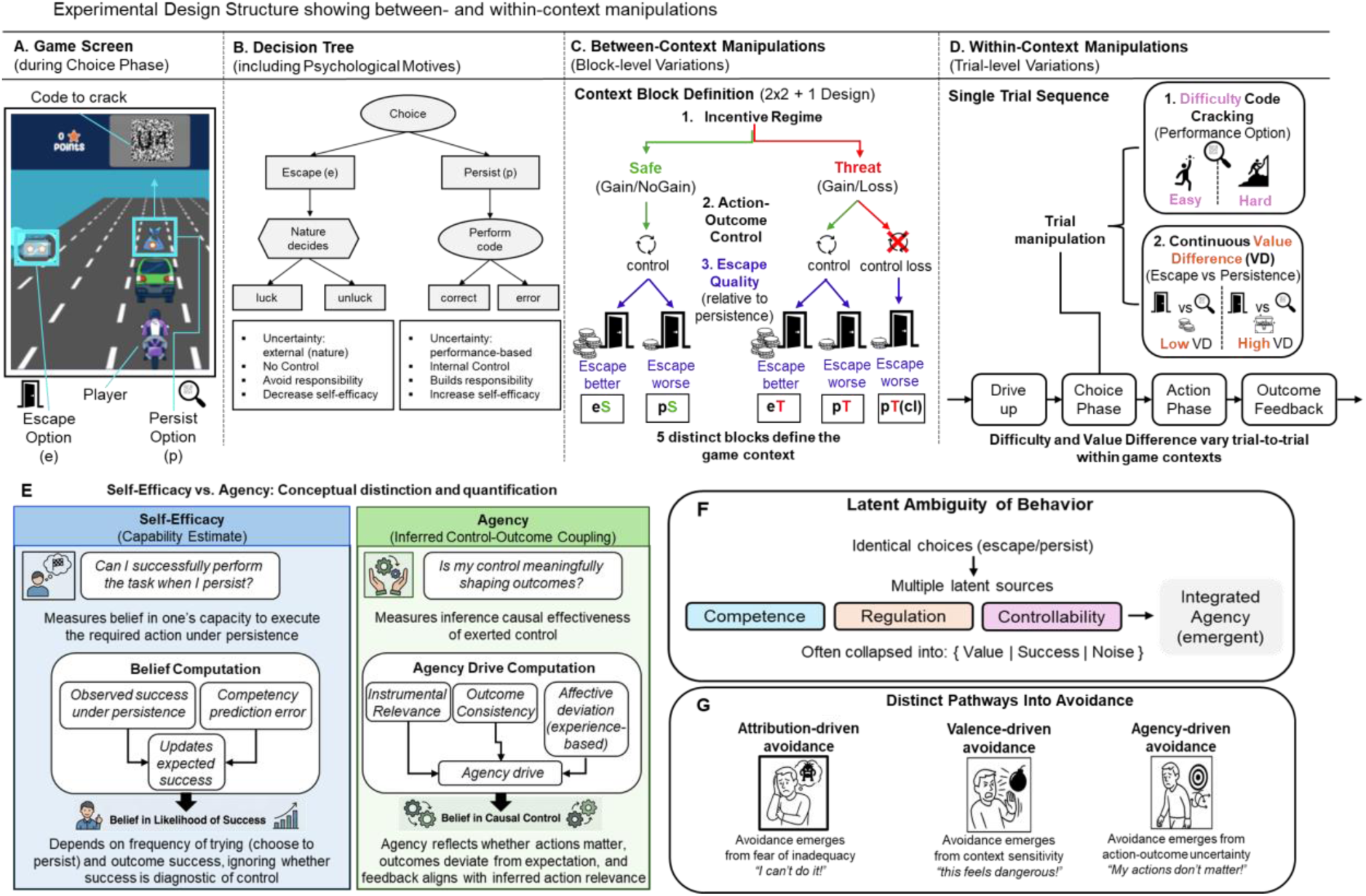
Overview of escape-persistence task structure and key avoidance constructs it captures. The paradigm dissociates *between-context policy arbitration* (adjusting persistence-escape strategies when the broader task architecture changes) from *within-context value sensitivity* (tracking graded changes in reward and effort without switching strategies). **(A)** On each *trial*, participants arbitrate between *persisting* in a goal-directed action and *escaping* via an explicit option. Persistence requires sustained effort under uncertainty (performing a code-cracking subtask) to achieve a goal (capturing a package). Escape allows disengagement at structured economic and psychological costs. **(B)** Schematic contrasting escape and persistence options that differ in source and control of uncertainty. Under escape, outcomes are determined by a nature-based process (luck vs. unluck), conferring no control over outcomes and potential psychological benefits (e.g., avoiding responsibility) alongside costs (e.g., reduced self-efficacy). Persistence outcomes depend on participants’ performance, granting greater control over outcomes but involving psychological costs (responsibility for failure). **(C)** Contextual factors, including incentive structure (safe vs. threat), relative advantage of escape (better vs. worse than persistence), and controllability over action-outcome mappings, are manipulated across blocks referred to as *game context*. **(D)** Reward magnitude and task difficulty vary trial by trial within game context (blocks). **(E)** Conceptual dissociation between self-efficacy and agency. *Self-efficacy* (left) reflects a belief about one’s capability to successfully execute an action, based on past success and expectations of performance. It tracks how likely one is to succeed when persisting, largely independent of whether success is causally driven by one’s actions. In contrast, *agency* (right) reflects a dynamically inferred belief about whether actions causally influence outcomes, integrating information about the instrumental relevance of actions, deviations from prior experience, and the consistency between expected and observed feedback, thereby determining whether outcomes are treated as informative for learning. This distinction allows for dissociations in which individuals feel capable of succeeding while simultaneously inferring that their actions do not matter. **(F)** The same choice can reflect different underlying processes. Controllability, competence, and regulation jointly affect *integrated agency*, but are often not dissociated. **(G)** Avoidance emerges via distinct latent pathways.

Identical escape patterns can arise from multiple latent factors (Fig. 1F) that can give rise to distinct computational pathways into avoidance (Fig. 1G). To dissociate them, our task incorporates several key features. First, embedding an independent calibration phase dissociates effort costs from feasibility. This allows a principled distinction between avoidance driven by effort aversion and avoidance driven by perceived control loss. Second, escape is implemented as a parametrically manipulable option whose advantage shifts across task contexts. This allows disengagement to be either adaptive or maladaptive depending on the context. Third, we separate policy-level adjustments from within-context value sensitivity by combining block-wise manipulations of incentive structure and controllability with trial-wise variation in reward and difficulty. These factors jointly contribute to an integrated estimate of agency. We define *agency* as a dynamically inferred quantity that reflects whether one’s actions are causally influencing outcomes (Fig 1E). Rather than a fixed trait, agency is continuously updated based on: 1. the inferred instrumental relevance of actions; 2. deviations between expected and experienced outcomes; and 3. the consistency between these signals, allowing attribution of outcomes to one’s own actions versus external factors. Accordingly, we conceptualize agency as a *gating signal* for learning that depends not only on inferred outcome controllability, but also on the reliability and coherence of the environmental feedback, thereby determining whether outcomes are treated as informative for learning. The task was implemented as an interactive, game-based mission within the *Gearshift Fellowship* research platform.^46^ This allows for task reconfiguration in an engaging and affectively salient setting.^15^

The primary study aim was to examine how escape behavior is flexibly calibrated to context. Rather than implementing a fully crossed factorial design of our new task structure, we configured the paradigm to differentially engage key drivers of avoidance (Fig 1C-D). This allowed us to test: 1. whether persistence under threat reflects the accumulation of prior agency-related evidence, rather than solely immediate responses to aversive cues; and 2. whether prior experience in contexts, where failure is safe and controllability can be learned, guides subsequent behavior under threat, consistent with a transfer of learned agency across contexts. To formalize how inferred self-efficacy and agency produce persistence versus escape decisions, we introduce the *Meta-Arbitration of Control and Agency Q-learning* (*MACA-Q; pronounced “MAH-kah-Q”)* model. It integrates instrumental and affective value signals within a unified framework and dynamically weights their interaction through meta-control processes that shape learning and arbitrate between competing decision strategies in a context-sensitive manner. MACA-Q allows us to: 1. simulate and characterize distinct pathways into avoidance; 2. quantify the rigidity of avoidance patterns; and 3. infer how beliefs about self-efficacy and agency emerge from the interplay of cognitive-affective evaluation processes.

The secondary study aim was to test whether dimensional variation in anxiety and depressive symptoms is associated with context-dependent patterns of avoidance. Our focus is not on clinical classification but on variation in symptom severity within a nonclinical sample. We therefore recruited a large online sample recruited via Prolific (N = 457), alongside self-report measures of anxiety and depressive symptoms (DASS-21).^47^

To preview our results, we find that avoidance behavior is not uniformly amplified under threat but depends critically on perceived controllability. When outcomes remained controllable, threat increased persistence. Consistent with this, high-performing individuals were not uniformly more persistent but more selective, escaping when persistence was no longer advantageous. We further identify distinct, context-dependent patterns associated with symptom dimensions that account for individual differences. Higher anxiety symptoms were associated with a disruption of the decision process itself. Specifically, individuals with higher scores showed reduced arbitration between effort and value and instead treated effort as a more unconditional cost. In this state, escape behavior remained responsive to heuristic cues but relatively insensitive to the incentive structure (i.e., whether failure occurs in a non-punitive context or is penalized). Conversely, higher depressive symptoms were associated with a distortion of the inputs to that process. This was reflected in a reduced ability to discriminate between effort costs and a pessimistic belief of task success under persistence.

Consistent with emerging theories of meta-control in neuroscience,^48–50^ and extending prior models that frame control as a resource to be allocated,^24^ MACA-Q conceptualizes meta-control as regulating the relative influence of value-based, heuristic, and affective signals in decision-making. We show that adaptive behavior is not explained by value learning alone but depends on meta-control processes that flexibly regulate decision strategies. This flexibility, grounded in dynamically inferred beliefs about agency and competence. This enables agents to sustain effort when control is meaningful and escape when it is not. Finally, model simulations show that prior experience in safe contexts can give rise to reduced avoidance under subsequent threat, consistent with the accumulation of agency-like beliefs across contexts.

## Results

We first characterize how persistence and escape vary across contexts. We then show that standard reinforcement learning models are insufficient to account for these patterns. To explain these dynamics, we introduce the MACA-Q model, which provides a mechanistic account of escape behavior under threat and dissociates distinct pathways into avoidance.

### Controllability under threat guides context-specific avoidance

Participants escaped more frequently when escape had a higher objective value than persistence [Mean escape frequency (MEF); MEF_escape better_ = 27.31%, MEF_persist better_ = 15.97%, *β* = −1.03, *SE* = 0.04, *p* < 0.001]. Holding constant the relative value difference between persistence-escape options, escapism differed systematically across incentive contexts. Participants escaped less under threat than under safe contexts [MEF_threat_ = 20.63%, MEF_safe_ = 24.30%, *β* = −0.29, *SE* = 0.04, *p* < 0.001]. This effect existed regardless of whether escape had a higher objective value (Fig 2A). This pattern differs from predictions of the classical avoidance-framing account, which suggest increased escape under threat.^51–53^

**Fig. 2.**
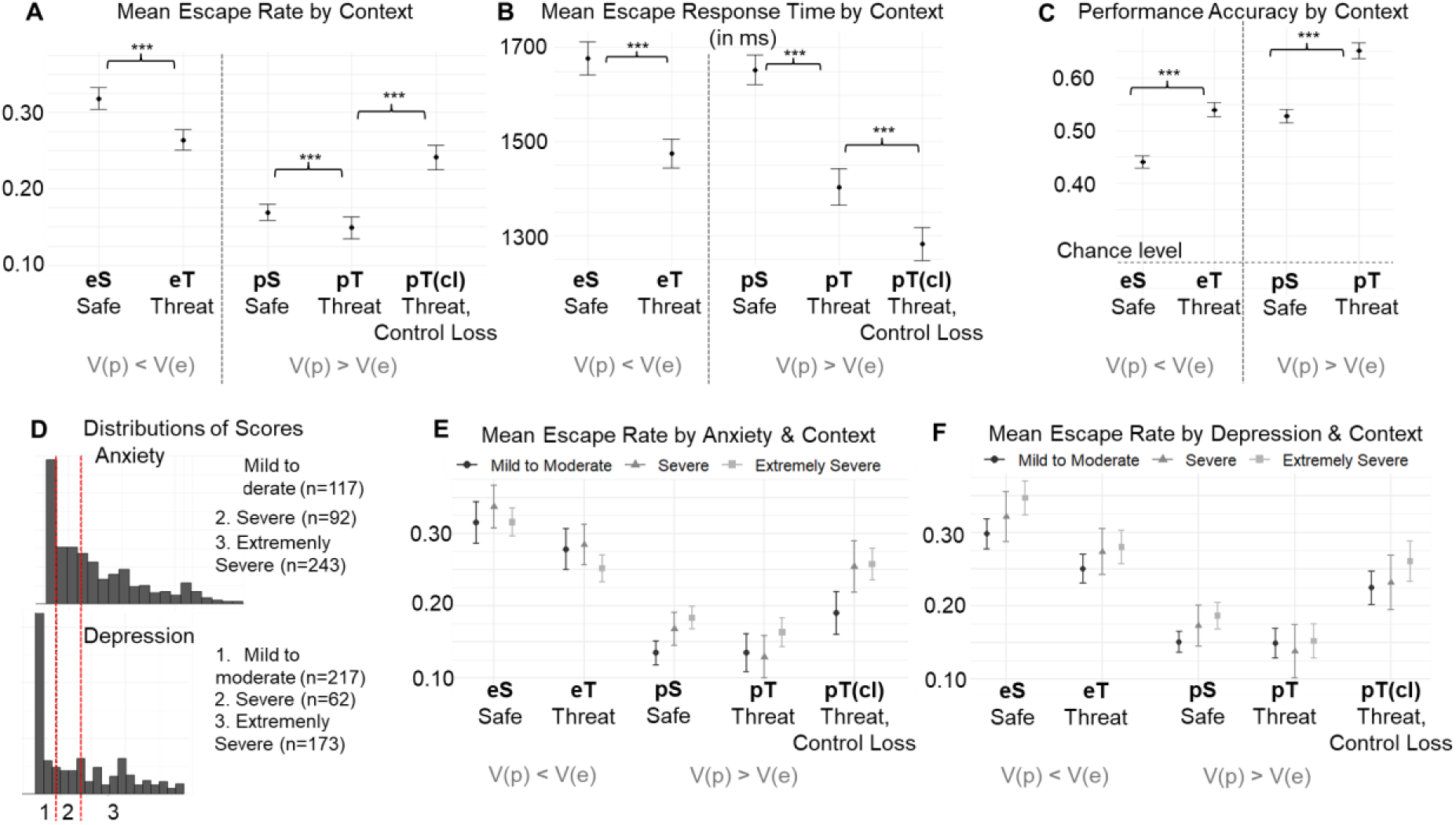
Behavioral escape pattern across context and clinical dimensions. The x-axis in the top-row panels (including E and F) corresponds to the block-level contextual manipulations introduced in Fig 1A. The first letter (e/p) indicates whether escape or persistence had the higher objective value (e: V(e) > V(p); p: V(e) < V(p)). The second letter (S/T) denotes the incentive regime, distinguishing safe contexts (gain vs. no gain) from threat contexts (gain vs. punishment). The pT_cl_ condition refers to a threat block in which persistence is objectively advantageous but controllability over outcomes is removed. For visualization, block contexts are shown in a fixed order (see for details Methods). The points denote means, which were calculated by first averaging within subjects and then across subjects. Vertical error bars indicate within-subject SEM. Asterisks indicate *p* < 0.001. **(A)** Mean escape rates were higher under safety than threat and increased when persistence no longer afforded control over outcomes. **(B)** Mean response times (RTs) were slower under safety than threat. Though, within threat contexts, RTs increased when escape was objectively more rewarding (eT vs. pT: *p* < 0.001), consistent with motivational conflict. **(C)** Under persistence, task accuracy was higher under threat than safety. Dashed line indicates chance level (25%) if subjects guess the correct answer. **(D)** Distribution of severity scores from the Anxiety and Depression subscales of the 21-item Depression Anxiety Stress Scales. **(E)** Mean escape rates by anxiety severity and context show higher escape rates under safety when escape was worse (pR) and under loss of control (pT_cl_). **(F)** Mean escape rates by depression score and context show higher escape rates under safety, but not threat, for highest severity.

A hallmark of our paradigm is that the persistence choice allowed participants to influence outcomes through their own effort (via task performance), whereas the escape option did not (Fig 1B). When this action-outcome contingency for choosing persistence was removed (control loss), escapism increased markedly, accounting for all other contextual factors (Fig 2A). Specifically, escape frequency increased by 10% relative to the comparable performance-contingent context [comparing pT vs pT_cl_ contexts: MEF_control_ = 14.89%, MEF_control-loss_ = 24.12%, *β* = 1.33, *SE* = 0.08, *p* < 0.001]. Hence, in the uncontrollable threat condition, escape patterns aligned with the behavioral predictions by the classical avoidance-framing account.

To further probe the decision dynamics underlying escapism, we examined response times (RTs) across incentive contexts (Fig 2B). Overall, escape decisions were faster under threat than under safe contexts, regardless of whether escape or persistence had a higher monetary value [Mean escape RT (MERT); MERT_escape_ = 1,500ms, MERT_persist_ = 1600ms, *β* = 0.08, *SE* = 0.02, *p* < 0.001]. Above and beyond this general speeding under threat, escape decisions under threat were significantly slower when escape was objectively more rewarding than persistence (comparing eT vs pT in Fig 2B; *β* = 0.16, *SE* = 0.03, *p* < 0.001). This suggests a selective decision slowing when the choice to escape involved a benefit-cost trade-off (i.e., maximizing monetary reward at expense to relinquishing outcome control).

We next examined how threat affected participants’ task performance under persistence (Fig 2C). Their accuracy was higher under threat than under safe contexts, despite identical task difficulty and relative value structure across contexts [Mean accuracy under safety = 59.46%, under threat = 65.85%, *β* = 0.41, *SE* = 0.04, *p* < 0.001]. This pattern suggests increased effort allocation for persistence under threat.

Because safety blocks always preceded threat blocks, cross-regime differences could, in principle, reflect time-on-task effects. However, the observed behavioral profile was not consistent with practice or fatigue effects. First, escape rates exhibited non-monotonic changes across blocks and increasing again in the last uncontrollable condition. Second, both escape behavior and performance accuracy under persistence tracked local contingency structure rather than elapsed time. Finally, all effects remained robust when controlling for trial-wise time-on-task in mixed-effects models (Supplementary Tables S3-S13).

### Individual differences in choice arbitration biases across contexts

Our online sample (N=457) spanned a broad range of anxiety and depressive symptom severity (Fig 2D), with distinct escape patterns across the three context-change dimensions (Fig 2E-F). We therefore tested whether individual differences along these dimensions modulated behavioral adaptation to contextual changes (i.e., escape advantage, incentives, and controllability). To this end, we fit a mixed-effects logistic regression model predicting trial-based escape choice from symptom scores and contextual factors (see Methods). Anxiety and depressive scores were moderately correlated (*r* = 0.64, *p* < 0.001) and did not exhibit problematic multicollinearity (*VIF* = 1.7). Both were therefore entered into the same model to estimate their unique contributions.

Higher depressive scores were associated with a higher escape probability across contexts [Fig 3A; *β* = 0.31, *SE* = 0.14, *p* = 0.030]. Higher anxiety scores were not associated with a global escape bias [*β* = −0.07, *SE* = 0.14, *p* = 0.587]. Critically, anxiety and depressive dimensions exhibited dissociable patterns of contextual sensitivity. Higher depressive scores were associated with increased value-based differentiation, reflected in stronger modulation of escape behavior by relative escape advantage (Fig 3B; symptom x escape value interaction: *β* = 0.26, *SE* = 0.06, *p* < 0.001). Conversely, higher anxiety scores were associated with increased escape even when it was monetarily better, leading to a negative escape value arbitration (Fig 3B; *β* = −0.39, *SE* = 0.06, *p* < 0.001). This suggests a reduced coupling between choice behavior and value signals. Anxiety scores were further linked to decreased incentive differentiation (Fig 3C; symptom x incentive interaction: *β* = −0.20, *SE* = 0.06, *p* < 0.001), driven by increased escape under threat [*β* = 0.02, *SE* = 0.01, *p* = 0.014]. This behavioral pattern is consistent with the hypothesis that anxiety drives threat overgeneralization.^42,44^ Conversely, higher depressive scores were linked to increased incentive differentiation [*β* = 0.20, *SE* = 0.05, *p* < 0.001], driven by increased escape under safety [*β* = 0.03, *SE* = 0.02, *p* = 0.051].

**Fig. 3.**
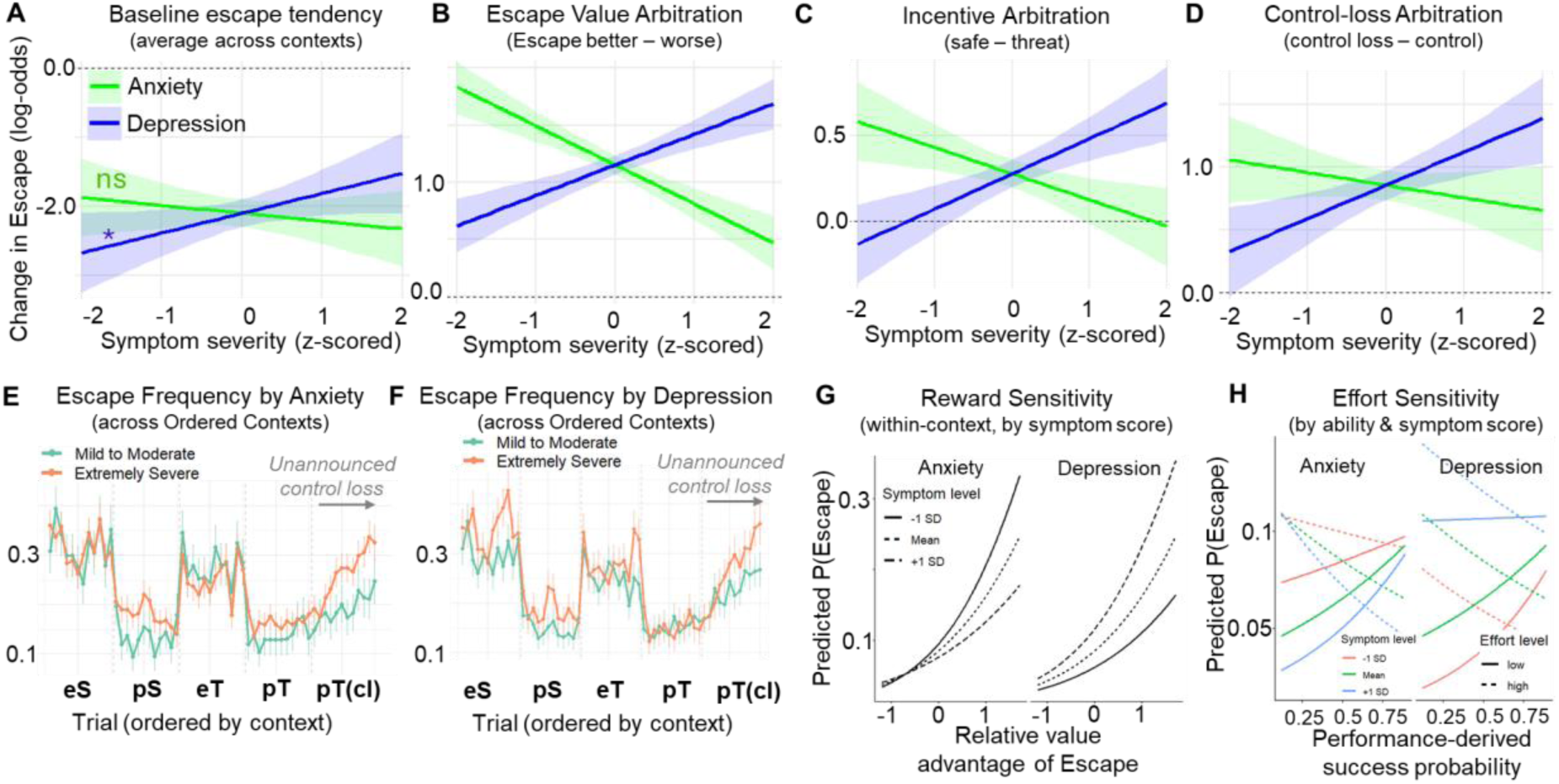
Predicted marginal effects of between-context arbitration and within-context sensitivity. The top row illustrates how escape probability (log-odds) varies with symptom severity (x-axis). **(A)** Baseline escape tendency averaged across contexts. The asterisk indicates a significant positive association between global escapism and depressive scores; no such association was observed for anxiety scores (ns). **(B-D)** Contrast-coded changes in escape probability between the two contexts being compared, with positive values indicating greater escape in the second context relative to the first. *Escape value* contrasts contexts in which escape was objectively better versus worse than persistence in terms of monetary reward. Negative slopes indicate value-consistent behavior, such that participants were less likely to escape when escape carried lower relative value, whereas positive slopes indicate value-insensitive escape, suggesting disengagement driven by factors other than relative reward. *Controllability* contrasts contexts in which outcomes under persistence were contingent on participants’ actions versus not. Effects were estimated using linear mixed-effects models predicting escape choice from symptom severity of anxiety and depressive symptoms and contextual contrasts. **(E)** Escape probability across trials as a function of anxiety scores, showing distinct updating of escape behavior following *unannounced* control loss. For visualization, block contexts are shown in a fixed order (see Introduction and Methods for details). Symptom severity was grouped into mild-to-moderate and extremely severe categories based on DASS-21 scores. **(F)** Same as (E), shown for depressive symptom severity, illustrating differential updating of escape behavior following unannounced loss of controllability. **(G)** Predicted escape probability as a function of reward sensitivity (i.e., the relative value difference between escape and persistence) shows that sensitivity was moderated by symptoms. Predictions are derived from linear mixed-effects models estimating escape choice as a function of reward sensitivity, contextual factors, task difficulty, ability-derived expected success (pSuccess), and (z-scored) symptom scores. For illustration, we show separate lines indicate symptom levels (−1 SD, mean, +1 SD). All other covariates are held constant (baseline block context, baseline effort, mean pSuccess). **(H)** Effort x pSuccess decomposition by symptom level (based on same model as in G). Predicted escape probability is shown with relative reward held constant at zero (no incentive advantage), isolating the interaction between nominal effort and pSuccess.

Increased escape under safety at higher depressive scores is consistent with prior work linking depression to escape-related biases.^54,55^ We extend this account by showing that this pattern reverses under threat, but only when persistence confers action-outcome control (Fig 3D; *β* = 0.45, *SE* = 0.21, *p* = 0.037]. Specifically, individuals with higher depressive scores showed reduced escape under threat when outcomes were controllable, relative to when they were not [*β* = −0.03, *SE* = 0.01, *p* = 0.006]. In contrast, higher anxiety scores were associated with reduced differentiation by controllability under threat [*β* = −0.24, *SE* = 0.22, *p* = 0.271]. This suggests a diminished use of control signals when arbitrating between persistence and escape. One alternative explanation is that the unannounced transition into the no-control context delayed the detection of contingency shifts in participants with elevated anxiety or depressive scores. However, participants with higher scores showed faster detection of control loss, as indicated by steeper escape trajectories following the transition (Fig 3E-F).

In sum, depressive and anxiety dimensions were associated with distinct behavioral adaptation modes. Higher depressive scores were linked to preserved, and in some cases enhanced, arbitration across contextual changes in value, incentive, and controllability. In contrast, higher anxiety scores were associated with escape behavior that was less sensitive to contextual structure, particularly under threat. Hence, symptom dimensions bias persistence-escape arbitration in a context-specific manner, rather than producing a global increase in avoidance.

### Individual differences in cost-benefit sensitivity within contexts

To isolate within-context valuation effects from both contextual policy shifts and individual differences in task ability, we modeled trial-wise escape choices as a function of block-level context, trial-level reward and effort, and an individual-specific estimate of success probability (pSuccess) assessed in a separate paradigm (see Methods). Effort indexed objective task difficulty (low vs. high), while pSuccess was derived from performance on the same task in an independent mission without escape alternatives. This allowed us to dissociate nominal task demands from participants’ objective likelihood of successful engagement (i.e., success likelihood under persistence separate from the strategic demands of this main paradigm).

As expected, higher reward magnitude for escape [*β* = 0.73, *SE* = 0.04, *p* < 0.001] and higher task effort [*β* = 0.15, *SE* = 0.06, *p* = 0.015] increased escape probability. However, escape decisions were not driven by capacity limitations per se, as pSuccess did not independently predict escape [*β* = −0.01, *SE* = 0.25, *p* = 0.950].

Individual differences in symptom dimensions selectively modulated benefit sensitivity. Higher anxiety scores were associated with reduced reward sensitivity for escape, controlling for context, effort, and success probability [Fig 3G; anxiety x reward interaction: *β* = −0.17, *SE* = 0.06, *p* = 0.002]. Conversely, higher depressive scores showed marginal reward sensitivity [Fig 3G; *β* = 0.10, *SE* = 0.05, *p* = 0.071]. Symptom scores did not modulate effort sensitivity (i.e., neither in terms of task difficulty nor pSuccess). Neither dimension significantly modulated effort sensitivity, whether indexed by objective difficulty or pSuccess. Together, these findings indicate that symptom-related differences in escape behavior primarily reflect altered benefit integration rather than effort sensitivity or capacity constraints. They motivate the subsequent examination of how effort costs are constructed from task competency.

### Individual differences in effort valuation: intrinsic cost & competency signal

Effort sensitivity has been proposed as a key mechanism in motivational dysfunction, yet most paradigms do not distinguish between objective difficulty and individual ability.^55–57^ It therefore remains unclear whether reduced engagement reflects an intrinsic aversion to effort or a belief that effort is unlikely to succeed.^18^ To dissociate these mechanisms, we extended the model to include the interaction between effort and competency (pSuccess), testing whether the subjective cost of effort depends on the likelihood of success.

Across participants, the effect of effort on escape was strongly modulated by competency [effort x pSuccess interaction: *β* = −1.692, *SE* = 0.22, *p* < 0.001]. Specifically, high effort increased escape primarily when success was unlikely, whereas its impact was attenuated when success was likely. Anxiety scores were associated with a global upregulation of effort costs [anxiety x effort interaction: *β* = 0.627, *SE* = 0.21, *p* = 0.002], independent of competency [Fig 3H; anxiety x effort x pSuccess interaction: *β* = 0.572, *SE* = 0.40, *p* = 0.153]. This pattern suggests that effort is treated as a context-insensitive cost signal, rather than being adaptively scaled by expected success. In contrast, depressive scores revealed a qualitatively different pattern. Higher scores were associated with reduced differentiation between low- and high-effort demands [depression x effort interaction: *β* = −0.687, *SE* = 0.20, *p* = 0.001]. However, when success was unlikely, depressive scores were associated with increased escape even for low-effort options [Fig 3H; depression x effort x pSuccess interaction: *β* = −0.937, *SE* = 0.37, *p* = 0.013]. This pattern suggests a loss of effort-based discrimination, coupled with a generalized bias toward escapism under low expected success.

Together, these behavioral patterns indicate that participants adjusted behavior in a context-sensitive manner, selectively engaging persistence when action-outcome contingencies were reliable and showing dissociable sensitivities to effort and prior success.

### Computational Mechanisms Underlying Adaptive Persistence

We next tested whether a canonical Q-learning model could account for the observed behavioral patterns, providing a baseline for evaluating the underlying learning processes.

### Standard reinforcement learning fails to capture key behavioral signatures

A canonical Q-learning model failed to capture key behavioral signatures (Fig 4 A-E, top row), which were captured by our extended *MACA-Q* model (Fig 4A-E, bottom row). We first explain where canonical Q-learning falls short, motivating the extensions in MACA-Q.

**Fig. 4.**
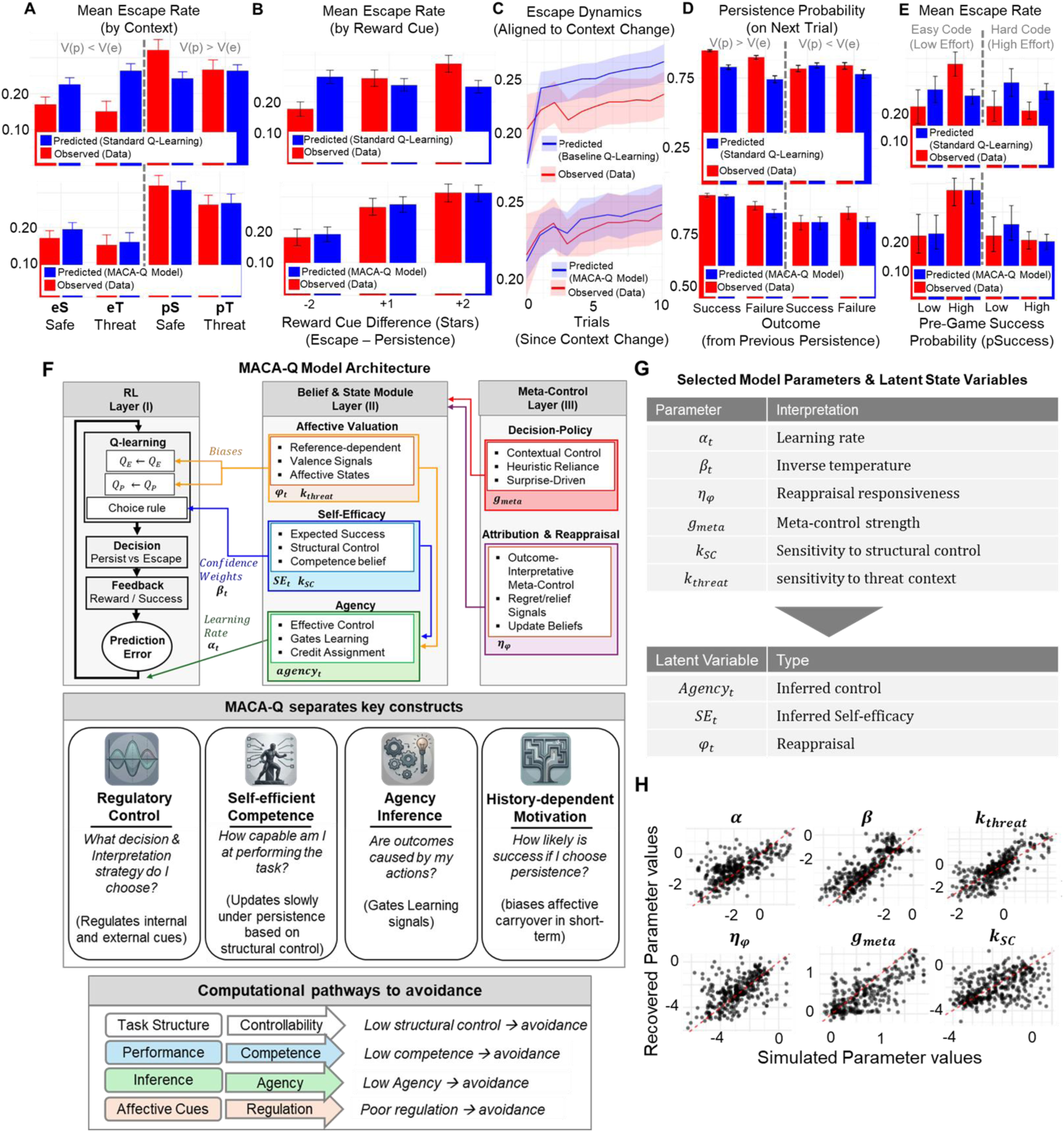
MACA-Q captures behavioral patterns beyond standard RL and dissociates key processes. (**A-E)** Model comparison of behavioral predictions (see Methods for details). Context labels: The first letter (e/p) indicates whether escape or persistence had the higher objective value (e: V(e) > V(p); p: V(e) < V(p)). The second letter (S/T) denotes the incentive regime, distinguishing safe contexts (gain vs. no gain) from threat contexts (gain vs. punishment). The pT_cl_ condition refers to a threat block in which persistence is objectively advantageous but controllability over outcomes is removed. For visualization, block contexts are shown in a fixed order. **Top:** Canonical Q-learning predictions (blue), which fail to capture key qualitative patterns in the observed data (red) across conditions. **Bottom:** MACA-Q model predictions reproduce these patterns, indicating that an integrated cognitive-affective architecture is required to account for persistence and escape behavior. (**F) Top:** Conceptual architecture of MACA-Q. A RL layer updates action values for persistence and escape based on experienced outcomes. These updates interact with control-inference modules that track beliefs about competence and controllability (self-efficacy and agency). A higher-level meta-control system regulates both outcome interpretation and decision policy: (re)appraisal processes shape how outcomes influence affective and efficacy states, while decision-level meta-control arbitrates between value-based and heuristic strategies. These processes operate across timescales, with fast affective responses, intermediate value updating, and slower accumulation of competence and controllability beliefs. **Bottom:** MACA-Q dissociates key cognitive-affective constructs, including regulatory control, self-efficacy, agency, and affective bias, that are typically conflated in standard RL models. **(G)** Overview of key model parameters and emergent latent variables, including agency, self-efficacy, and appraisal states. Latent variables arise from interactions among model parameters and are computed from the model’s internal dynamics rather than directly estimated. **(H)** Results from parameter recovery analyses (see Methods). Parameters were sampled from the empirical distribution, used to simulate behavior, and re-estimated. Recovered parameters closely matched ground truth (red dashed line: identity), indicating reliable parameter identifiability (all r > 0.5; see also Table S5 & Fig S1).

First, empirical escape rates were lower under threat than safety contexts, irrespective of whether escape was more or less advantageous (Fig 4A, top: red bars). The canonical Q-learning model, relying on a stationary environment, predicts the opposite pattern (Fig 4A, top: blue bars). This is because increasing the penalty associated with failed persistence should reduce its expected value and promote escape. Second, escape rates were modulated by contextual cues signaling the relative advantage of escape over persistence (Fig 4B, top: red bars). This suggests that participants’ choices was also guided by heuristic cues (reward stars) beyond incrementally learned values. Canonical Q-learning predicts that choices depend solely on learned action values. Third, adaptation following context changes unfolded gradually (Fig 4C, top: red curve). this indicates that participants did not immediately switch strategies but instead updated them incrementally, consistent with uncertainty about the new context contingencies. Canonical Q-learning predicts rapid adjustment via prediction-error-driven updating as it lacks an explicit representation of uncertainty about context contingencies. Fourth, outcome-based updating depended on task structure (Fig 4D). Failures reduced persistence only when persistence was the preferable option, but not when escape was advantageous. Canonical Q-learning predicts uniform updating following negative outcomes. Finally, prior achievement (pSuccess) modulated escapism selectively rather than globally (Fig 4E). Higher-achieving participants escaped more when persistence offered little value but converged with others when persistence was clearly advantageous. Canonical Q-learning predicts a uniform shift in choice bias. Together, these empirical patterns indicate that choices were guided by hierarchical control processes that integrate learning and inference processes.

### The Meta-Arbitration of Control and Agency Q-Learning Model (MACA-Q)

MACA-Q extends standard Q-learning by embedding value-based choices within a hierarchical system that infers controllability, tracks internal states, and arbitrates control allocation. The architecture includes three interacting layers (Fig 4G-F): At the RL layer I, the model maintains separately learned action values for persistence and escape which are updated via prediction-error learning (*α_t_*) similar to the canonical Q-learning model.

Unlike standard RL though, these updates are not applied uniformly but are gated by belief & state modules in the second layer which maintain three latent variables: 1. Affective valuation process (*φ_t_*) gates the influence of contextual biases (e.g., threat sensitivity, *k_threat_*) onto learned action values. By integrating prediction errors relative to a long-term affective baseline, it allows the model to become more sensitive to feedback under threat (rapidly updating decision biases independently of reward history). 2. Self-efficacy process (*SE_t_*) integrates expected success likelihood (*pSuccess_t_*) and structural control sensitivity (*k*_SC_) to accumulate a latent competency belief. This belief dynamically tunes decision precision (*β_t_*), such that high efficacy promotes consistent goal commitment while low efficacy increases choice stochasticity (escapism). 3. Agency dynamics (*agency_t_*) acts as a congruence gate for credit assignment. Rather than tracking objective success alone, it computes the alignment between internal competency expectations and external feedback. This ensures that prediction errors only update action values when outcomes are perceived as contingent on the agent’s strategy. In so doing, it formalizes how a loss of agency can suppress learning even in the presence of rewards.

At the third level, meta-control processes arbitrate the relative importance of second-layer processes. First, they regulate outcome interpretation through (re)appraisal mechanisms (*η_φ_*), attributing unexpected outcomes to either noise or structural shifts. Second, they also regulate the decision policy via a meta-control gain (*g_meta_*), which determines the relative weighting of learned values versus heuristic strategies. By separating value learning from higher-order inferences, MACA-Q provides a unified account of how persistence and escapism emerge as context-sensitive outcomes of interacting cognitive and affective processes. See Methods for details and full mathematical formalization.

### MACA-Q captures behavioral pattern and provides identifiable parameters

We evaluated MACA-Q using three complementary validation analyses (see Methods). First, predictive checks showed that the model accurately reproduced the empirical behavioral signatures (Fig 4A-E, bottom panels). Second, parameter recovery analyses demonstrated reliable identifiability of key parameters (Fig 4; Fig S1). Third, systematic ablation analyses revealed that removing individual components selectively impaired the model’s ability to capture specific behavioral signatures (Fig S2-4). Hence, each component contributes uniquely to model performance. Formal model comparison further confirmed that the MACA-Q model version introduced here outperformed simpler model versions (Table S16).

### Behavioral equivalence masks computational trajectories

As outlined in the Introduction, similar behavior can arise from distinct underlying mechanisms and only become dissociable across contexts. We therefore identified parameterizations of learning rate and meta-control that produced comparable escape rates in some contexts (Fig 5A). Despite similar behavior, these parameterizations differed in their underlying agency dynamics (Fig 5B) and gave rise to distinct escape patterns in subsequent threat contexts, where behavior is particularly sensitive to agency (Fig 5C). Hence, similar behavior can arise from distinct latent computational trajectories that only become dissociable across contexts.

**Fig. 5.**
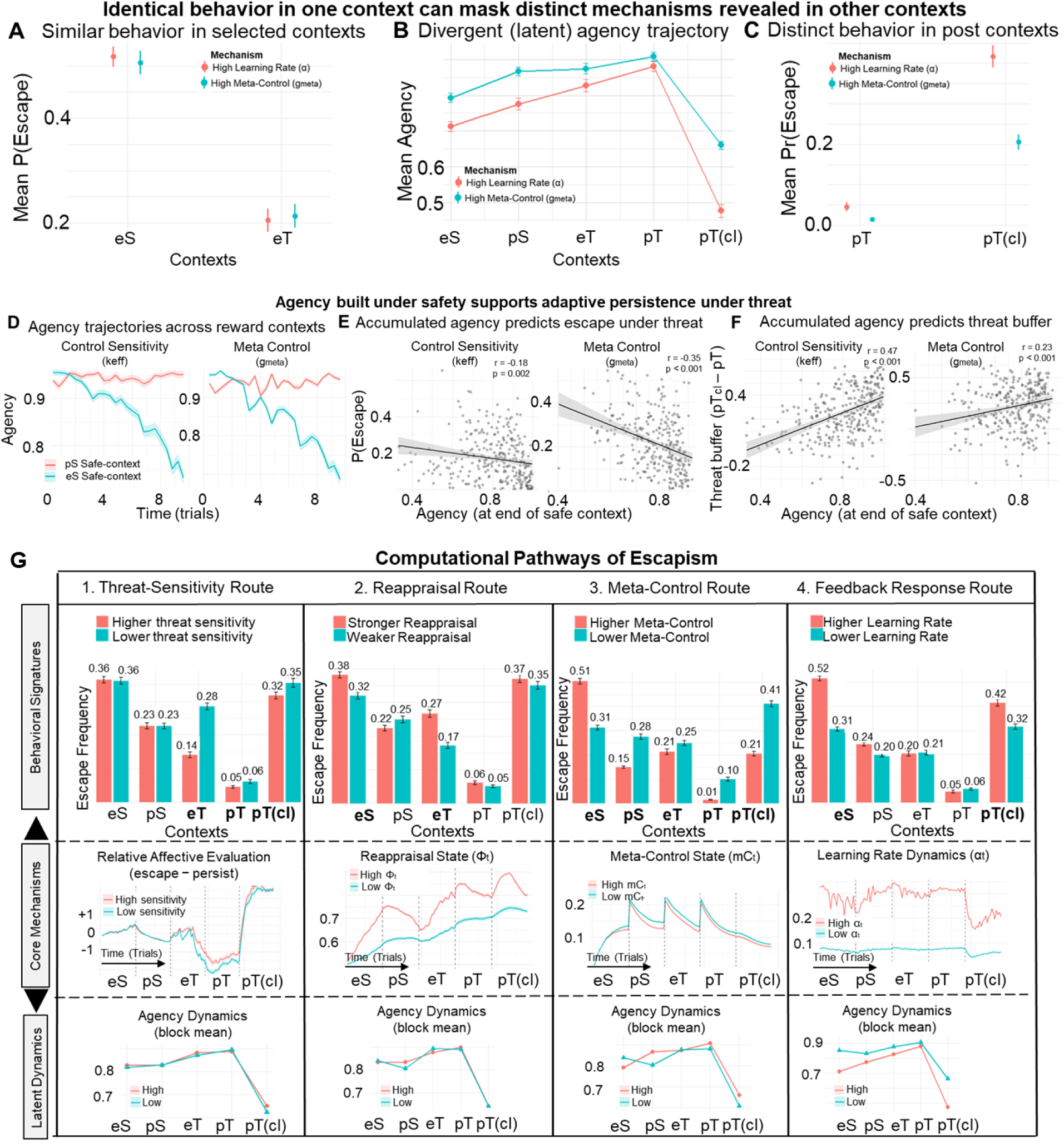
Cross-context dissociation of mechanisms and agency dynamics. **(A-C):** Identical behavior in a single context can arise from distinct computational mechanisms that are only revealed across contexts (see Fig. 1C for abbreviations). **(A)** Empirical escape rates under safety and threat when escape is advantageous (eS, eT). Error bars indicate 95% confidence intervals across simulated subjects (n=400). **(B)** Mean inferred agency across contexts under high learning rate versus high meta-control. The meta-control pathway shows higher and more stable agency, whereas the learning-rate pathway exhibits lower agency and a sharper decline under reduced controllability. **(C)** In diagnostic threat contexts with and without control (pT, pT_cl_), the same model variants produce divergent escape behavior, revealing mechanistic differences not apparent in focal contexts. Error bars indicate 95% confidence intervals. **(D-F)** Agency built under safety predicts adaptive persistence under threat through distinct computational mechanisms. **(D)** Trial-wise trajectories of inferred agency during safety contexts (pS: persistence advantageous; eS: escape advantageous). Despite identical external contingencies, parameter manipulations produce distinct agency dynamics. **(E)** Across 400 simulated individuals, higher agency at the end of safety contexts predicts reduced escape under subsequent threat when persistence remains advantageous (pT). Each point represents a simulated subject; lines denote linear fits. **(F)** Agency built under safety predicts a threat-buffering index: difference in escape between uncontrollable (pT_cl_) and controllable (pT) threat. Higher agency supports selective disengagement: persistence is maintained when control is preserved and escape increases when control is lost. **(G)** Simulation-based dissociation of computational pathways. Isolated manipulation of key parameters reveals distinct routes to escape behavior (top row). The behavioral signature of each route is highlighted by bolding the corresponding context on the x-axis. For each pathway, varying a single parameter within its empirical range (middle row) produces characteristic latent dynamics and corresponding changes in inferred agency (bottom row). Increased threat sensitivity biases behavior toward persistence under threat, reappraisal promotes adaptive switching when persistence becomes suboptimal, higher meta-control stabilizes agency and supports flexible adjustment, and higher learning rates increase sensitivity to recent outcomes, leading to faster escapism and reduced agency.

### Accumulated agency is associated with reduced escape under threat

Building on the hypothesized protective role of agency (see Introduction), we tested whether agency accumulated under safe contexts, where failure under persistence is safe, relates to subsequent escape under threat. We focused on meta-control, which guides goal-directed decision consistency and structural control sensitivity as key drivers of agency formation. Across simulations, higher agency at the end of safe contexts was linked to reduced escape under subsequent threat (Fig 5D-E). This effect was contingent on controllability (Fig 5F). When outcomes remained action-contingent (pT), higher agency reduced escape. When control over threat was removed (pT_cl_), higher agency increased escape. This demonstrates that accumulated agency does not merely promote blind persistence. Instead, it enables adaptive differentiation between controllable and uncontrollable threat. Together, this suggests that agency provides a unifying mechanism linking distinct computational pathways to behavior.

### Distinct Computational Pathways into Escapism

MACA-Q dissociates multiple mechanisms that are typically conflated in models of avoidance and reward learning, including affective valuation, outcome interpretation, meta-control, and learning dynamics (Fig 5G). To characterize these pathways, we varied (in isolation) key parameters governing threat sensitivity (*k_threat_*), reappraisal attunement (*η_φ_*), meta-control strength (*gₘₑₜₐ*), and feedback responsiveness (*α*); see details in Methods.

#### Threat-sensitivity route

Variation in trait-level threat sensitivity determines how strongly threat biases affective valuation. Higher threat sensitivity reduced escape under threat when persistence remained controllable by amplifying the affective contrast between persistence and escape (Fig 5G: column 1, top panel). This increased the relative affective value of persistence, promoting engagement under threat without altering learning dynamics in other contexts. Because this bias operates at the decision policy level, it does not alter inferred agency trajectories (Fig 5G: column 1, bottom panel). Hence, affective states can flexibly bias decision-making under uncertainty independently of learning or controllability inference, consistent with prior work.^58–60^

#### Reappraisal route

Variation in reappraisal responsiveness determines how rapidly outcome-related appraisal states (e.g., regret, relief) are updated. Stronger reappraisal responsiveness increased escape when it was advantageous by accelerating the downregulation of persistence following worse-than-expected outcomes. This effect operates through outcome interpretation relative to contextual expectations. It selectively engages at points of increased appraisal demand, including contextual boundaries, threat onset, and control loss (Fig 5G: second column, middle panel). Since this pathway primarily modulates outcome interpretation, variation in reappraisal responsiveness had only modest effects on inferred agency.

#### Meta-control route

Variation in meta-control determines how strongly contextual information is integrated into the decision policy. Higher meta-control led to more adaptive behavior across contexts (Fig 5G: column 3, top panel), with more escape when it was advantageous and less when it was not. This reflects enhanced alignment of behavior with the current value structure, enabling rapid adjustment across contexts. Conversely to the threat-sensitive and reappraisal pathways, meta-control also influenced the evolution of inferred agency. Higher meta-control supported stable accumulation of agency through consistent engagement with persistence under safety. Instead, lower meta-control led to more variable behavior and noisier agency trajectories. These findings indicate that meta-control not only governs adaptive switching but also stabilizes the accumulation of agency signals that support flexible behavior across contexts.

#### Feedback responsiveness route

Variation in learning rate determines how strongly value estimates are updated in response to feedback. Higher learning rates increased escape under safe contexts where escape was advantageous and under conditions of control loss by making value estimates more reactive to recent outcomes. When persistence outcomes deteriorate, the value of persistence declines more rapidly, promoting faster escapism. In contrast to the threat-sensitive and reappraisal pathways, variation in learning rate also strongly influenced inferred agency. Namely, higher learning rates produced lower and more volatile agency estimates, whereas lower learning rates preserved more stable value estimates and sustained a sense of control. During unannounced control loss, higher learning rates led to faster declines in agency. Conversely to meta-control, which stabilizes behavior and promotes the accumulation of agency through consistent engagement, higher learning rates amplify reactivity to recent outcomes, resulting in lower agency accumulation and reduced stability of inferred control.

Together, these findings establish that persistence and escape emerge from distinct computational pathways that are organized through dynamically inferred agency, providing a unifying framework for understanding how avoidance arise across contexts.

## Discussion

Deciding when to hold on and persist versus when to let go and avoid are fundamental to adaptive behavior under uncertainty and partial outcome control. Avoidance has often been studied as a response to threat or cost.^6–8^ In many real-world settings though, these factors are intertwined with beliefs about controllability and competence, as well as with affective demands (e.g., threat and outcome framing). These factors jointly guide how outcomes are experienced and regulated. This makes it difficult to disentangle the underlying mental processes that give rise to avoidance. We developed a paradigm that systematically varies the aforementioned contextual factors and administered it to a large online sample (N=457). We showed that this provides a complementary perspective to the classical accounts of avoidance. Specifically, persisting versus escaping emerge from a hierarchical arbitration decision process that integrates incentive structure, inferred controllability, cognitive-affective signals and prior experience across contexts. To formalize this process, we introduced the MACA-Q model. Unlike standard RL, which could not account for the observed behavior, MACA-Q embeds value-based choice within a meta-control architecture that regulates how outcomes are interpreted and used for learning and decision-making. In this model, agency emerges as a dynamically inferred quantity, rather than a fixed trait. Notably, we resolve a central ambiguity in the study of avoidance by integrating cognitive-affective theories of control,^24,55^ meta-control,^48–50^ and emotion regulation^58,61–64^ into an extended Q-learning framework. Specifically, we show how similar behavioral patterns can arise from distinct underlying mechanisms that remain difficult to dissociate when examined within a single context. Using our paradigm and model simulations, we show that agency accumulates in contexts where failure is safe and actions reliably predict control. This accumulated agency then carries over to subsequent threat contexts, where it buffers against disengagement when persistence is advantageous.

To examine how escape behavior is calibrated by context, our task structure formalizes avoidance as an explicit choice between persistence and escape under varying conditions. Unlike standard approach-avoidance paradigms,^8,15,19,33–35,65^ escape is implemented as a parametrically manipulable option whose advantage shifts across contexts. This allows escapism to be either adaptive or maladaptive depending on task context. Combining block-wise manipulations of incentive structure and controllability with trial-wise variation in reward and task difficulty, the design separates policy-level adjustments from within-context value sensitivity. This allows to dissociate mechanisms that are typically conflated, linking effort valuation, competence beliefs, and controllability inference within a unified framework.^15,46^ Lastly, the task combines explicit heuristic cues with latent variables that must be inferred through experience, allowing us to dissociate value-based learning from higher-order inferences about controllability and agency.

Our findings reveal a central role for effort in adaptive behavior. Rather than functioning as a primitive aversive cost,^18^ we found that effort acts as an informational signal that is dynamically interpreted in relation to expected competence. When expected success is high, the subjective cost of effort is attenuated. This promotes sustained engagement even under threat. Conversely, when expectations of control are low, the same effort signals lead to escapism. This view aligns with recent accounts suggesting that effort does not merely reflect cost.^3,66^ Instead, it also functions as a signal of value that contributes to meaning construction and thereby potentially also to the inference of agency. We integrate this view by formalizing agency as an emergent property of cognitive-affective evaluation processes rather than from value learning alone.

We found that high-performing individuals were not uniformly more persistent but more selective, escaping when persistence was no longer advantageous. These results indicate that adaptive persistence reflects the efficient regulation of engagement based on inferred controllability and competence, rather than a simple willingness to exert effort. Recent work suggests that reduced flexibility in updating decision strategies and beliefs are transdiagnostic, underexplored clinical characteristics.^15,67^ Consistent with this view, our analyses of individual differences indicate that clinical symptom dimensions disrupt persistence-escape arbitration in distinct ways. These characteristics become observable at a behavioral level when examining avoidance across contexts. Higher anxiety severity was associated with a reduced ability to utilize accumulated competence as a buffer. As such, effort was treated as a relatively fixed cost regardless of prior success. This pattern suggests a shift toward cost-dominant evaluation, leading to escapism that is less sensitive to contextual incentives. In contrast, depressive symptoms were characterized by reduced differentiation between effort levels and reduced success expectations. This weakened the influence of actual competency on behavior. Together, these results show that avoidance is not a unitary construct but emerges through distinct computational pathways, with implications for understanding heterogeneity in maladaptive behavior.

Across findings, behavior could not be explained by self-efficacy or effort costs alone because participants did not uniformly disengage under threat or task difficulty. Instead, their persistence depended on whether outcomes were inferred to be controllable. Building on this, we formalize agency as a computational quantity distinct from self-efficacy (Fig 1D). Their dynamic interaction then determines when persistence versus disengagement is adaptive. The MACA-Q model formalizes this process, capturing how they are learned from different signals and integrated over time. This dissociation explains why individuals may feel capable yet disengage when outcomes become uncontrollable, or persist under threat when control is preserved. This is captured in our model simulations, showing that experiences in failure-safe contexts reduce subsequent maladaptive avoidance (i.e., avoidance when persistence is objectively better) under threat. Hence, what generalizes is not merely competence, but a belief about the causal efficacy of one’s actions. Together, our results indicate that adaptive behavior is guided by the joint inference of capability and controllability, formalized here as distinct but interacting constructs of self-efficacy and agency.

Conceptually, MACA-Q implements a hierarchical three-layer architecture: an *Actor* for value learning, an *Observer* that functions as a “*composer*” by actively constructing meaning through the integration of affective valuation, self-efficacy, and agency signals, and a *Governor* for meta-control. Rather than passively monitoring states, this intermediate layer composes task-relevant meaning by integrating affective, efficacy, and agency signals. This structure captures how affective states and reappraisal mechanisms dynamically guide instrumental engagement and learning. These hierarchical control dynamics relate also to existing computational accounts of cortico-striatal learning architectures.^68–74^ In particular, models of prefrontal-basal ganglia (PFC-BG) circuits propose layered decision systems in which higher-level contextual representations regulate downstream action-selection policies.^75,76^ Within such architectures, prefrontal signals gate the expression of behavioral strategies rather than merely updating action values through prediction errors.^77–79^ The meta-control component formalizes this principle by introducing a higher-level arbitration signal that modulates the relative influence of competing decision components, such as effortful persistence and lower-effort escape policies, based on contextual cues and ongoing experience.

By integrating value learning, control allocation, affective valuation, and motivational dynamics, MACA-Q connects several lines of research that have been studied in isolation. Specifically, RL accounts emphasize how action values are updated through experience, whereas cognitive-control theories focus on how control is allocated based on expected costs and benefits.^24^ Affective models highlight how expectation-dependent states guide valuation and bias choice,^58,80^ and motivational theories emphasize how effort is expended in pursuit of reward.^3,81^ MACA-Q provides a unified account of how persistence and escapism emerge from the interaction of learning, control, and affective processes.^12,45^ It formalizes meta-control as an arbitration over the relative influence of value-based, heuristic, and affective signals, determining when persistence remains worthwhile. This perspective is consistent with theories of control allocation and extends them by incorporating affective valuation and outcome reinterpretation as integral components of decision-making.^48–50^ It also aligns with psychological accounts emphasizing the role of competence beliefs and perceived controllability in regulating motivation.^14,82–86^ Compared to these frameworks, we leverage their insights to formalize how agency and self-efficacy emerge as distinct but interacting computational quantities that jointly guide adaptive persistence versus escape.

Our simulations further clarify the regulatory mechanisms that enable flexible calibration of persistence and escape. Meta-control operates as a higher-order process that orchestrates the influence of contextual cues and heuristic signals on decision policies. By stabilizing engagement with controllable actions, it indirectly shapes inferences about agency. This view is consistent with accounts proposing that cognitive control systems arbitrate between competing strategies based on environmental demands.^87,88^ The architecture also incorporates affective valuation, allowing emotional signals to track reward history and bias choice.^58,80,89^ In MACA-Q, affect is not epiphenomenal but interacts with competence beliefs and controllability inferences to shape valuation and behavior. A key component of this system is a reappraisal pathway, which enables reinterpretation of outcomes. For example, failure can be reframed as a signal of task difficulty rather than lack of ability, thereby reshaping the relative value of persistence and escape. By embedding reappraisal within the learning architecture, MACA-Q provides a mechanistic account of how emotional reinterpretation supports adaptive flexibility and links RL and hierarchical control^24^ with theories of self-efficacy^85,86^ and learned helplessness.^12,45^ In so doing, MACA-Q builds and extends from related models of hopelessness and escape behavior that distinguish controllability from belief dynamics.^90^ Compared to existing models, they do not implement reappraisal as an explicit regulatory pathway within a trial-wise learning-and-decision architecture, nor do they formalize the distinction between self-efficacy and agency.

While reappraisal is central to cognitive-behavioral theories of emotion regulation, it has only recently been incorporated into computational models of decision-making.^61–64,87,88,91^ These existing approaches remain largely separated along three distinct lines of work. One line of work models the selection of regulation strategies, such as when individuals choose reappraisal versus distraction as a function of emotional intensity and temporal trade-offs.^92^ These models formalize strategy choice but do not address how regulation (re)constructs the meaning of outcomes within ongoing instrumental learning. A second line of work, often grounded in active-inference frameworks, models how cognitive restructuring and exposure update beliefs about threat and safety.^91,93^ These approaches make CBT-style belief updating computationally explicit, but primarily focus on changes in threat-related beliefs. They do not explicitly dissociate self-efficacy from agency, nor do they model agency as a latent quantity that accumulates across contexts and shapes subsequent persistence under threat. A third line of work in computational psychiatry isolates how specific intervention components affect distinct mechanisms, such as effort sensitivity or attributional style.^62,64^ While this provides important mechanistic insights, these models do not embed regulatory processes within the same architecture that governs value learning, control allocation, and persistence-escape decisions.

Compared to this existing work, MACA-Q is distinctive in combining these elements within a single framework. It embeds reappraisal-like reinterpretation within an ongoing arbitration architecture that jointly updates value, inferred controllability, self-efficacy, and agency. These latent states in turn determine whether persistence remains meaningful in a changing environment. This integration allows the model to capture how regulatory processes produce not only beliefs, but also guide adaptive decisions across contexts and over time.

Rather than mapping individual parameters onto symptom categories, we leverage MACA-Q as a generative computational model to characterize latent dynamics that unfold over time. Parameters determine how strongly a system responds, whereas latent states reveal when specific processes are engaged by the environment. A central implication of these analyses is that resilience emerges from prior experience through the accumulation of agency. Adaptive behavior under threat is therefore not a fixed tendency to persist, but the capacity to regulate persistence based on inferred controllability. Individuals with higher agency do not persist indiscriminately; they maintain engagement when outcomes remain controllable and escape when control is lost. This reframes resilience as the flexible regulation of persistence rather than persistence itself. Our results further suggest that adaptive behavior emerges from the capacity to infer when control is lost in a specific context and to act on that inference. Persistence in the absence of controllability constitutes a failure of arbitration, not a virtue. Instead, adaptive agents disengage and reallocate effort toward contexts where their actions remain consequential. Avoidance, in this sense, is not inherently maladaptive. Instead, it becomes maladaptive only when it generalizes beyond the contexts in which control is truly diminished.

The implications of our work extend to neuroscience, economics, artificial intelligence, and psychiatry. In neuroscience, the model generates testable predictions about how hierarchical cortico-striatal circuits support adaptive behavior. Specifically, how prefrontal systems track inferred controllability and regulate the influence of value-based and affective signals on action selection. By linking latent computational variables such as agency and meta-control to distinct functional timescales, the framework offers a principled approach to identifying neural signals that govern the flexible regulation of engagement. In economics, disengagement emerges as a rational response to reduced agency rather than a failure of motivation. This has direct implications for organizational design and leadership in environments that demand continuous adaptation. Adaptive behavior is not characterized by indiscriminate persistence or avoidance, but by the selective reallocation of engagement based on perceived controllability. When individuals infer that their actions no longer meaningfully influence outcomes, or when action-outcome contingencies become unreliable, continued investment becomes maladaptive. Model simulations and empirical results show that the most adaptive systems disengage and redirect effort toward contexts in which control can be effectively exerted. From this perspective, avoidance is not a global bias, but a context-sensitive policy that preserves agency by reallocating resources toward controllable opportunities. In artificial intelligence, the framework provides a blueprint for agents that can autonomously decide when to persist in effortful computation based on environmental feedback. Extending recent proposals that agents must represent what aspects of the environment matter for decision-making,^94,95^ and building on work on rational meta-reasoning and control allocation,^48,83^ our results suggest that adaptive agents must also infer whether their actions causally influence outcomes. Incorporating such agency inference would enable agents to disengage from uncontrollable contexts and reallocate resources toward environments where effective control can be exerted. In psychiatry, it enables the decomposition of behavioral phenotypes into mechanistic pathways, supporting a dimensional and transdiagnostic approach. Rather than mapping single parameters to symptom dimensions, we use the model to identify specific breaks in a universal control architecture. This approach may provide quantifiable markers for cognitive-behavioral interventions, such as Acceptance and Commitment Therapy and related mindfulness-based approaches, that aim to modify how individuals respond to challenge, uncertainty, and distress.^96–100^

Several limitations should be considered and addressed in future research. First, incentive regimes were not counterbalanced, as safe contexts preceded threat contexts to capture how experience affects agency and subsequent escape behavior. Although the observed patterns are not consistent with simple practice or fatigue effects, future work using counterbalanced designs will be necessary to isolate causal effects of incentive structure. Second, while the MACA-Q model was validated through predictive checks, ablations, and parameter recovery, further work is needed to link its latent variables to neural and physiological measures. Especially parameters of slower meta-learning processes showed weaker recovery and were therefore not emphasized in the main text (see Supplement). This likely reflects a limitation in the temporal depth of the current paradigm, as such processes unfold over longer timescales. Future work using extended gameplay or longitudinal designs will be important to more reliably capture these dynamics. Finally, symptom effects were assessed in a nonclinical sample, and future work should examine whether these computational signatures generalize to clinical populations. A key direction is to characterize how distinct disruptions in controllability inference and meta-control arbitration give rise to symptom dimensions associated with anxiety and depression. Anxiety may arise from a bias toward overestimating the probability or cost of control loss, lowering the threshold for disengagement. In contrast, depression may involve impairments in updating self-efficacy and agency following negative outcomes, resulting in persistent under-engagement even when control is reinstated. Beyond disengagement, a critical but underexplored process concerns the re-emergence of engagement. Namely, how agents recover from failed persistence and re-enter goal-directed behavior. Computationally, this requires modeling how agency is reconstructed, how beliefs about controllability are revised over time, and how meta-control mechanisms re-enable effort allocation. Characterizing these recovery dynamics may provide a mechanistic account of resilience and its disruption in affective disorders.

The present work illustrates the potential of reconfigurable, game-based missions to probe adaptive behavior in dynamic environments. The paradigm was implemented within the Gearshift Fellowship research platform,^46^ which provides a naturalistic framework for studying decision-making in affectively salient and continuously evolving contexts. This approach moves beyond static behavioral summaries toward dynamic, mechanism-sensitive assays. Recent studies have already advocated for such an approach as it enables the study of how individuals learn when to persist, when to disengage, and how these decisions evolve across environments.^15,34,46,67^ More broadly, such frameworks open the possibility of interactive systems that not only measure latent cognitive and affective processes, but also reflect them back to the individual, supporting the development of more adaptive interpretations and decision policies over time.^15,46,101^

## Data availability

Data are available in the manuscript and supplementary materials. It will be made available via OSF at the publication stage.

## Materials and methods

### Participants

A total of 500 participants were recruited through the online research platform Prolific.^102^ Recruitment used Prolific’s built-in prescreening filters, which allow researchers to target participants based on demographic and self-reported characteristics recorded in their Prolific profiles. All procedures were approved by the Brown University Institutional Review Board, and all participants provided informed consent prior to participation.

Participants were required to be between 18 and 35 years of age and currently residing in the United States. Recruitment was further stratified using Prolific prescreening criteria such that approximately half of the sample self-reported having received a diagnosis of a mental health condition, whereas the other half reported no prior mental health diagnosis. Recruitment was stratified to achieve approximately equal numbers of male and female participants. Participants were required to complete the study in a single uninterrupted session using a computer desktop and standard keyboard.

After providing informed consent, participants first completed a battery of self-report questionnaires hosted on Qualtrics. For the present study, the primary measure of interest was the Anxiety and Depression subscales of the 21-item Depression Anxiety Stress Scales (DASS-21; see subsection *Questionnaire*).^47^ Afterwards, participants were automatically redirected to the Gearshift Fellowship research platform,^46^ where they completed the experimental game paradigm described next. The entire session, including instructions, questionnaires, and gameplay, lasted approximately 60 min. Participants received a fixed payment of 15 USD at the end of study completion.

### Experimental Paradigm

Our paradigm can be understood as an experimentally controlled persistence-escape arbitration *task* that embeds opportunities for disengagement alongside sustained, effortful goal-directed action. The paradigm was implemented within the Gearshift Fellowship digital research platform.^46^ Completing the game took approximately 30 minutes. There was a total of 60 trials (12 per block).

### Game flow

On each trial, participants chose between a persistence option and an escape option. Choosing the persistence option allowed participants to earn the reward by completing a cognitively demanding code-cracking task. Because outcomes following persistence depended on participants’ task performance, persistence represented a partially controllable pathway for obtaining rewards (Fig 1A). In contrast, the escape option allowed participants to bypass task performance but at the cost of relinquishing control over the outcome (Fig 1B). This is because outcomes following escape were determined externally rather than by participant performance.

After choosing to persist or escape, participants steered their driver into the corresponding lane. Selecting the persistence option required engaging in the effortful code-cracking task to obtain the reward (see below), whereas the escape option yielded the alternative outcome without effort. Importantly, after trial-wise feedback, the driver was always returned to the center, such that the persistence option was directly ahead and thus constituted the default option (Fig 1A). We therefore refer to choosing the target option with the code-cracking task as *persist*, as it requires continued engagement with the task.

### Code-cracking during persistence

If participants selected the persistence option, they entered a code-cracking task that required identifying the correct response based on perceptually degraded stimuli presented in the gray window above the target car (Fig 1A). The target car cued which feature of the stimulus (letter or number) was relevant. Participants then made a categorical decision based on the cued dimension: for letters, they indicated whether the stimulus was a consonant or vowel; for numbers, whether it was odd or even. Participants were informed prior to the task which target car corresponded to each stimulus dimension, ensuring that the cue-dimension mapping was unambiguous.

Stimulus degradation varied across trials to manipulate task difficulty and the amount of cognitive effort required. For trials in which the letter dimension was relevant, low-difficulty stimuli were generated using a background brightness of 45%, number brightness of 10%, and letter brightness of 90%, resulting in high perceptual contrast for the letter. High-difficulty trials retained the same background brightness but reversed the brightness levels of the number and letter (number brightness = 90%, letter brightness = 10%), thereby substantially reducing the perceptual contrast of the letter relative to the distractor dimension. For trials in which the number dimension was relevant, low-difficulty stimuli used a background brightness of 55%, number brightness of 10%, and letter brightness of 90%, producing high contrast for the number. In high-difficulty number trials, the brightness values were adjusted to reduce the perceptual contrast of the number relative to the letter while keeping the background brightness constant. Similar stimulus manipulations have been employed in previous studies, where their psychometric properties (i.e., their effects on perceptual encoding difficulty and task performance) have been characterized.^103^

### Trial-based Feedback

Following the decision (and task execution if persistence was selected), participants received outcome feedback. They steered their driver into the corresponding lane and observed whether the package landed in that lane. Feedback was presented visually as a package dropping onto a conveyor line, indicating whether the action had succeeded. In the persistence condition, successful completion of the code-cracking task yielded the reward associated with that trial, whereas incorrect responses resulted in no reward. In the escape condition, outcomes were determined by the externally imposed risk structure signaled during the choice phase.

### Heuristic cues of cost-benefit trade-offs within contexts

The potential reward associated with each option (persistence versus escape) was signaled through a visual cue. Specifically, the number of stars displayed on each package (1-3 stars; see Fig 1A). These cues indicated low, medium, or high reward prospects. Importantly, the exact reward values associated with each cue were not explicitly revealed, requiring participants to learn the expected value of each option through trial-by-trial experience. Specifically, one-star packages were associated with rewards drawn from a uniform distribution with a mean of 2 points (SD = 2), two-star packages from a distribution with a mean of 5 points (SD = 2), and three-star packages from a distribution with a mean of 8 points (SD = 2). On each trial, the realized reward was randomly sampled from the corresponding distribution and presented as numerical feedback above the selected package. Participants were not informed about these distributions. Thus, although reward magnitude was cued, participants still needed to learn expected values from stochastic outcomes. Critically, adaptive decisions further depended on additional task dimensions not directly specified by these cues, including success probability, effort costs, and outcome controllability.

The potential risk associated with the escape option was signaled through a visual heuristic cue. Namely, a frontbus appearing on the horizon and moving down the screen in a separate lane. This cue indicated the likelihood that choosing the escape option would result in a negative outcome (i.e., no reward in the safe context or a punishment in the threat context which we detail next). When no frontbus was present, the escape option carried no risk. A yellow frontbus indicated a low probability of punishment (0.20), whereas a red frontbus indicated a high probability (0.80). These outcomes were externally determined and not under the participant’s control, such that escape outcomes were probabilistic and uncontrollable. Reward-risk trade-offs were counterbalanced across block types (contexts), as detailed below. As the present study focuses on controllability and persistence, and high-risk trials were relatively sparse, analyses of risk were not a primary focus.

Participants were informed about the meaning of the visual cues prior to the start of the paradigm (i.e., that stars indicated potential reward magnitude and frontbus cues indicated the probability of risk). However, the precise reward values and outcome probabilities associated with these cues were not disclosed and had to be learned over the course of the experiment through trial-by-trial feedback.

### Between-block (context) manipulations

The paradigm comprised five block types: eS, eT, pS, pT, and pT_cl_. Each block included 12 trials. The first letter (e or p) indicates whether escape (e) had a higher objective value than persistence (p), V(e) > V(p), or vice versa V(e) < V(p). The second letter denotes the incentive regime (Safety [S] or Threat [T]). In the safe context, successful actions yielded points, whereas incorrect responses under persistence (code-cracking task) or capturing a bomb during escape resulted in zero points. In the threat context, the same events resulted in the loss of the points that could have been earned on that trial. Thus, identical behavioral outcomes produced either neutral (zero) or negative (loss) consequences depending on incentive context. The final block (pT_cl_) introduced an unannounced degradation of controllability under threat, such that correct actions no longer reliably determined outcomes.

Incentive regimes were presented in a fixed order, with safe blocks (eS, pS) preceding threat blocks (eT, pT, pT_cl_). This design avoided exposing participants to an aversive loss context at paradigm onset, consistent with prior evidence that incentive order influences behavior.^104^ Within each incentive regime (eS vs pS; eT vs pT), block order was counterbalanced across participants (between-subjects). Contextual changes (in incentive structure or escape advantage) were signaled by the number of stars displayed on the choice packages, allowing participants to infer the current context. However, these contexts were not explicitly labeled a priori. Moreover, the loss of controllability in pT_cl_ was not signaled by any explicit cue. Instead, participants had to infer this change from mismatches between their actions and the resulting outcomes. By combining these manipulations, the paradigm dissociated factors that are typically confounded in avoidance paradigms, including cognitive effort, incentive valence, externally imposed threat, and perceived controllability. Participants were thus required to continuously evaluate whether persistence remained worthwhile relative to escape.

### Questionnaire

Anxiety and depressive symptoms were assessed using the Anxiety and Depression subscales of the 21-item Depression Anxiety Stress Scales (DASS-21).^47^ The DASS-21 provides dimensional indices of symptom severity in nonclinical and community samples and does not constitute a diagnostic instrument. Standard convention was followed by multiplying subscale scores by 2 to enable comparability with the original DASS-42 cut-offs. For descriptive purposes, symptom severity categories were collapsed to reduce sparsity (Mild and Moderate combined), with no participants falling into the Mild category alone. All analyses therefore treat anxiety and depressive symptoms as continuous, dimensional variables rather than indicators of categorical clinical diagnoses.

### Computing individual-specific global competence (pSuccess)

Our experimental paradigm was embedded within a multi-mission interactive game as part of the Gearshift Fellowship platform.^46^ This enabled us to use participants’ performance from an independent calibration mission to estimate each participant’s probability of success under persistence (pSuccess). The calibration task was administered prior to the main paradigm. It consisted solely of the code-cracking component of the persistence option, without additional strategic elements (e.g., escape choices or contextual manipulations). The code-cracking component used the identical stimulus structures from our paradigm. For each participant, pSuccess was computed as the overall accuracy across all trials in the calibration mission. Importantly, this estimate was aggregated across difficulty levels to capture a participant-specific, task-general measure of competence. This approach allows us to dissociate stable success expectations (pSuccess) from context-dependent effort allocation in the main paradigm, which is driven by trial-wise trade-offs between task difficulty and reward under varying contexts (block types).

By estimating pSuccess from an independent task without an escape alternative, we avoid circularity between performance and choice. This ensures that competence is not conflated with strategic disengagement. The participant-specific measures were then used in the main analysis to approximate initial expected success under persistence. This allows the model to distinguish between limitations in ability and context-dependent inference of control. Moreover, this separation ensures that variation in effort expenditure is not trivially absorbed into competence estimates, allowing us to isolate how individuals respond to increasing task demands.

### Statistical Analyses

Of the 500 recruited participants, 43 did not complete the study due to premature termination (n=3), technical issues (n=10), or withdrawal of consent prior to beginning the experiment (n=30). The final dataset therefore included 457 participants, which were included in the subsequent analyses. All descriptive and statistical analyses were conducted in R and for linear mixed models, we used the established lme4 package.^105^

To characterize descriptive differences across contexts (block types), we analyzed mean escape rate (MER) and response times (RTs) using linear mixed-effects models with block type as a fixed effect and subject-specific random intercepts. RTs were log-transformed prior to analysis to account for the positive skewness typical in RT distributions.^35,106^ MER was computed at the subject-by-block level, allowing the use of linear mixed models for summary-level comparisons. A random-intercept structure was used to account for repeated measures while maintaining model stability and interpretability for descriptive comparisons. These analyses were intended to provide descriptive confirmation of behavioral differences across contexts.

The primary statistical analysis focused on three contrasts based on the three task manipulations (Fig 1C): incentive framing (safe vs. threat contexts), relative expected value (escape-favorable vs. escape-unfavorable contexts), and outcome controllability under persistence (present vs. absent). These contrasts were evaluated as predefined linear combinations of estimates and model-implied predictions derived from a single fitted model (described below). This allowed us to test targeted hypotheses about context-dependent arbitration of escape behavior rather than global escape tendencies, while grounding all inferences in a common parameterization. To this end, trial-wise escape choices were analyzed using a generalized linear mixed-effects model (logistic link), including a subject-specific random intercept to account for repeated measures. The model included a categorical predictor for block type, representing the five task conditions that jointly encode incentive context, relative expected value of escape versus persistence, and action-outcome controllability. Continuous measures of anxiety and depressive symptoms (z-scored) were included as predictors, along with their interactions with block type. Contrasts were computed post hoc from this model as linear combinations of block-level estimates. As a sensitivity analysis, we additionally fit an equivalent model in which these task dimensions were entered directly as contrast-coded predictors. Results were consistent across parameterizations, confirming that inferences did not depend on the model specification (Supplementary Table S8 & S9). All continuous predictors were mean-centered, and model estimates are reported on the log-odds scale. For visualization, model-implied effects were computed by simulating from the fixed-effect covariance matrix and evaluating predicted contrasts across the observed range of clinical symptom severity.

To control for multiple comparisons within the primary hypothesis space, we applied false discovery rate (FDR) correction across the set of predefined symptom-by-contrast interaction tests. Results were not materially altered by FDR correction, and the pattern of significant and non-significant effects remained unchanged. In the main text, we report unadjusted p-values from the primary regression models. The supplementary Table S2 presents both uncorrected and FDR-adjusted results.

To assess the potential influence of task order, practice, and fatigue effects arising from the fixed sequencing of incentive regimes (safety preceding threat), we conducted additional sensitivity analyses including a trial-wise time-on-task covariate (global trial index) in the mixed-effects models. This covariate captures monotonic changes in behavior over the course of the experiment, independent of task context. All primary models were re-estimated with this additional predictor, and the key effects of interest remained qualitatively and statistically unchanged (Supplementary Tables S3-S9). These analyses indicate that the observed context-dependent differences in escape behavior cannot be explained by global time-on-task effects.

The secondary statistical analysis focused on within-context sensitivity to parametric task variables, namely: reward sensitivity (i.e., the effect of changes in the relative monetary value between the escape and persistence options) and effort sensitivity (i.e., the effect of changes in task difficulty). To this end, we extended the primary trial-level generalized linear mixed-effects model to examine whether symptom variation was associated not only with context-dependent shifts across block types, but also with sensitivity to these parametric task variables within blocks. Specifically, we added predictors indexing nominal task difficulty (*effort*) and the relative value advantage of escape over persistence (*rewardSens*), together with their interactions with anxiety and depressive symptoms (z-scored). These models tested whether symptom dimensions were associated with altered weighting of effort costs or escape-related value differences beyond the categorical structure captured by block type. In these models, we additionally included a subject-level estimate of task ability under persistence (*pSuccess*) as a covariate. Including pSuccess allowed us to assess whether symptom-related differences in escape behavior remained associated with effort and reward sensitivity after accounting for individual differences in task ability. In a further extension, we included the interaction between effort and pSuccess, as well as its interaction with symptom dimensions, to test whether the effect of nominal task difficulty depended on objective feasibility and whether this dependency varied as a function of symptoms. All secondary models used the same outcome variable (trial-wise escape choice), logistic link function, and subject-specific random intercept structure as the primary model.

### Computational Analysis: Q-Learning Baseline Model

Statistical analyses characterize how task manipulations influence behavior but do not formalize the underlying generative processes that give rise to these effects. To complement this approach and provide a mechanistic account of decision-making, we implemented computational models grounded in cognitive-affective theory. As a baseline comparison to our MACA-Q model (introduced in the next section), we first implemented a canonical Q-learning model in which action values for persistence (Target) and escape (Escape) were updated via reward prediction errors. Specifically, on each trial t, action values (a) were updated as follows:

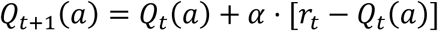

Separate learning rates were used for positive and negative prediction errors to allow for asymmetric updating from gains and losses. Choices were generated using a softmax decision rule with inverse temperature parameter β, such that the probability of choosing *Escape* was given by:

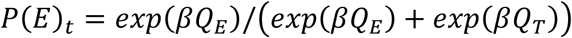

The model was applied to the same task structure as the empirical experiment, including trial-wise safe and threat contexts, block transitions, and probabilistic outcome contingencies. Action values for both options were initialized to zero and updated only based on experienced outcomes, without incorporating higher-order representations of controllability, agency, or context.

Model parameters (learning rates and inverse temperature) were estimated for each participant using the same fitting procedure as the MACA-Q model (see Method subsection: Computational Fitting Procedure). Model performance was evaluated based on its ability to reproduce key behavioral signatures, including context-dependent escape rates and adaptation across block transitions (see Method subsection: Model Validation and Robustness Checks). As shown in Fig 4A-E (top rows), this canonical Q-learning model failed to account for the key empirical patterns which suggest that decision-making processes arise from a hierarchical arbitration of distinct contextual and valuation signals.

### Computational Analysis: The MACA-Q model

Choice data was modeled using the Meta-Arbitration of Control and Agency Q-learning (MACA-Q) model. MACA-Q extends the Q-learning baseline model by formalizing decision-making as a hierarchical process in which value learning is embedded within a broader system that infers controllability, tracks internal states, and arbitrates the allocation of control. It incorporates mechanisms that allow: (1) context-sensitive modulation of behavior beyond learned values; (2) inference over latent task structure; and (3) selective updating of action values contingent on inferred controllability and expected outcomes. By separating value learning from higher-order inferences about controllability, competence, and context, MACA-Q captures how agents decide whether exerting control is worthwhile. All variables evolve trial-by-trial and are formally specified below. Fig 4g summarizes the main parameters and Supplementary Table S14 provides a complete list of parameter and state variables. A summary of estimated parameters can be found at the end of this section.

### Trial-based inputs and observed variables

On each trial t, the agent observes reward outcomes (r_t_), effort costs (c_t_), and contextual cues such as dStars_t_ denoting the value difference between the options and the risk_t_ (indicated by the frontbus). The agent chooses between persistence (P) and escape (E) options.

### Reinforcement learning updating

The model maintains separate action values for persistence (*Q_P,t_*) and escape (*Q_E,t_*). The initial persistence value is derived from participants’ baseline success probability (*pSuccess*) by transforming it onto a latent value scale. The escape value is initialized at zero. Action values are updated via prediction errors. For escape, learning follows a standard Q-learning rule:

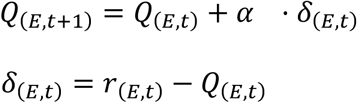

For persistence, learning is selectively gated by an inferred agency signal (detailed below):

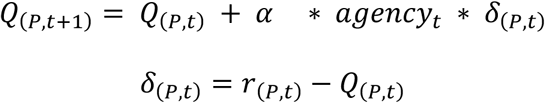

This asymmetry reflects the core assumption that outcomes only update persistence values when they are interpreted as contingent on one’s actions. In both equations, the term α denotes the standard learning rate.

### Action Q values and choice rule

On each trial, action values are integrated with affective signals, effort costs, and contextual biases to form latent utilities.

For persistence:

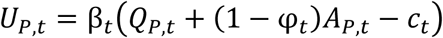

For escape:

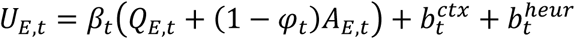

Here, β_t_ denotes the inverse temperature term. A_P,t_ and A_E,t_ denote option-specific affective signals, c_t_ denotes effort costs, and *φ*_*t*_ represents the current level of reappraisal. The gating term (1 − *φ*_*t*_) implements dynamic regulation of affective influence, enabling flexible reweighting of affect relative to instrumental value. The term *b_t_^ctx^* denotes slower contextual biases, and *b_t_^heur^* denotes dynamic within-context heuristic biases favoring escape. Note that contextual biases occur outside the softmax temperature: β_t_ * Q + bias. Moreover, utilities are modeled as linear combinations of value, cost, and affective components, providing a parsimonious approximation that allows independent contributions of each process.

Effort costs entered the persistence utility term as a function of both trial difficulty and expected success. Specifically, effort costs were applied only on high-effort trials and were scaled by the complement of the current expected success belief, such that:

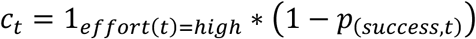

Thus, effort becomes more costly when tasks are difficult and expected success is low. Choice probabilities are given by a logistic decision rule:

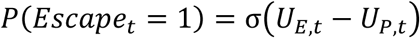

where σ(⋅) is the logistic function. This logistic rule formally characterizes the decision as a binary arbitration between two distinct behavioral strategies: persistence (goal-directed effort) and escape (strategic disengagement). Unlike a standard multi-action softmax, which treats all options as competing for the same resource, the logistic link function effectively models the probability of selecting the escape option relative to the currently maintained persistence policy.

### Self-Efficacy Module: Competency buildup across timescales

MACA-Q distinguishes three related but non-identical constructs that operate at different timescales. First, the model tracks an expected probability that persistence will succeed, *p_success,t_*, which is updated only on persistence trials through a success prediction error:

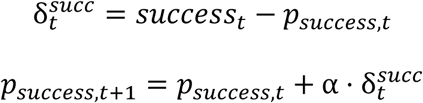

Where success_t_ ∈ {0,1}. This state captures evolving beliefs about the feasibility of successful persistence in the current environment.

Second, on each trial the model computes an instantaneous efficacy signal, *SE_inst,t_*, indexing the current structural advantage of persistence. In the implementation, this signal is defined as a scaled difference between the current persistence value and an expected value under a random choice policy (*EQ_TRP_*) which serves as baseline reference point:

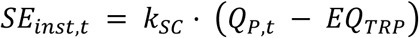

where *k_SC_* controls sensitivity to perceived structural control. This trial-wise signal reflects how advantageous persistence currently appears relative to a weak-control baseline.

Third, a slower latent competence state integrates efficacy-related experience signals over time:

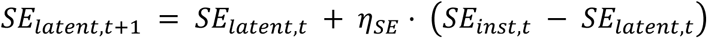

This state is updated primarily following persistence trials and only after post-choice appraisal and endorsement processes determine how strongly the current trial should be consolidated into the broader self-evaluative belief state. Escape choices do not provide direct evidence for one’s ability to succeed through persistence; accordingly, when escape is chosen, the latent competence state relaxes back toward neutrality rather than being reinforced. Hence, the term η_SE_ reflects an integration rate of self-efficacy that controls the updating of beliefs about action success probability based on recent performance.

The latent competence further modulates choice precision:

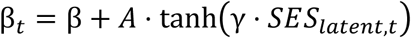

Functionally, this mechanism allows repeated evidence of effective control to shift behavior toward more stable exploitation of persistence when warranted, whereas weak or unstable competence beliefs maintain a flatter decision policy. This feature gives MACA-Q a computational interpretation of self-efficacy as a gain-control process: self-evaluative beliefs do not merely summarize past performance, but regulate how strongly the agent commits to the currently favored action. The terms A and γ control the strength and nonlinearity of modulation and were fixed at constant values. The hyperbolic tangent function ensures smooth, bounded modulation of decision precision, preventing unrealistically large fluctuations while preserving sensitivity to changes in self-efficacy. An additive formulation was chosen to allow self-efficacy to modulate, rather than replace, baseline decision stochasticity.

### Affective Module: Reference-dependent valuation and carry-over memory

In addition to value learning, the model includes a separate affective valuation system that tracks the emotional significance of outcomes relative to recent experience. Specifically, the system evaluates outcomes against a dynamic affective reference point (*R̅_t_*), which represents a running estimate of recent reward history. After each trial, the reference point is updated toward the experienced outcome according to:

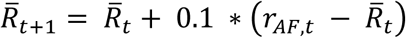

where *r_AF,t_* denotes the affective representation of the observed outcome. In the present model specification, the transformation linking raw outcomes to affective outcomes was fixed at 0.1.

Affective valence was defined as the deviation of the experienced outcome from this reference point. Specifically, the model computed a normalized valence signal:

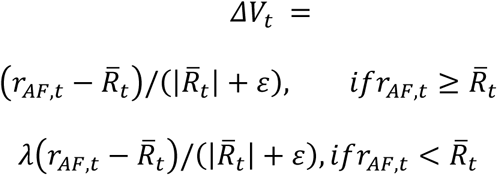

where λ > 1 increases sensitivity to worse-than-expected outcomes. This formulation implements reference-dependent affective evaluation with asymmetric weighting of negative deviations. The loss-weighting factor λ was fixed to 1.5 and ɛ was fixed to 0.01. This reflects the well-established asymmetry whereby losses exert greater impact than equivalent gains, consistent with empirical estimates from prospect theory.^107^

Separate affective states are maintained for persistence and escape:

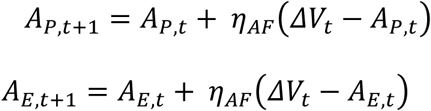

This allowed affective consequences of persistence and disengagement to accumulate differently over time. These option-specific affective states then biased subsequent choice by shifting the perceived utility of persistence and escape in the decision rule as shown before (see subsection: *Action Q values and choice rule*). In the winning model specification, affective updating was additionally modulated by threat context. When outcomes occurred in threat conditions, the effective affective learning rate (*η_AF_*) increased, making the emotional state more sensitive to deviations from the reference expectation.

This formulation corresponds to an exponential moving average, allowing the model to track recent affective outcomes while retaining memory of prior experience. This choice balances stability and adaptability and is widely used in reinforcement learning models of affective valuation.

### Agency Module: Inferred control and credit assignment

A central feature of MACA-Q is that learning from persistence outcomes is gated by a trial-wise inferred agency signal, *agency_t_*, which quantifies whether recent evidence supports the interpretation that persistence exerts a causal influence on outcomes. Agency is computed from three components: (1) the inferred instrumental relevance of persistence relative to baseline (*SE_inst_*), (2) the affective deviation of outcomes relative to an experience-dependent reference point (*ΔV_t_*), and (3) the consistency between inferred action relevance and experienced feedback.

Consistency between expected and experienced outcomes is defined as:

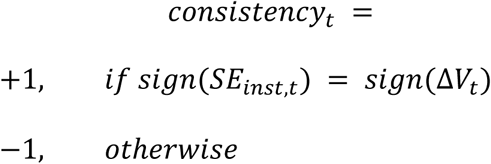

Formally, agency is defined as a bounded latent variable:

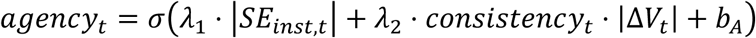

where σ denotes the logistic sigmoid function and b_A_ is a basleine bias term that was fixed to 0. The sigmoid function ensures that agency is bounded in the interval [0,1], consistent with an interpretation as a probabilistic belief over controllability. This also introduces nonlinear sensitivity, such that moderate evidence produces gradual changes, while strong consistent evidence leads to rapid consolidation of perceived control. This formulation is related to Bayesian accounts of controllability inference, where agency reflects the inferred probability that outcomes are contingent on actions, but is implemented here in a tractable parametric form.

The multiplicative interaction between consistency_t_ and *ΔV_t_* ensures that confirmatory evidence strengthens agency, whereas contradictory evidence weakens it.The contribution of each component is weighted by its magnitude, such that stronger efficacy signals and larger deviations from expectation exert a greater influence on agency. The parameters *λ*_1_, *λ*_2,_ were fixed a priori and were not estimated during model fitting.

The resulting agency estimate multiplicatively scales the persistence learning rate (α). Consequently, identical persistence outcomes can produce different value updates depending on whether they are interpreted as informative about controllable performance versus stochastic or uncontrollable variation. Following escape choices, the escape value is updated from its outcome, whereas the agency signal is not recomputed from instrumental relevance, affective deviation, and consistency. Instead, agency decays multiplicatively, reflecting that escape provides no direct new evidence about whether persistence is causally effective.

### Attribution and reappraisal mechanisms

MACA-Q includes a post-choice regulatory process that determines how strongly recent outcomes influence subsequent decisions. This process is captured by a dynamic reappraisal state, *φ*_*t*_ ∈ [0,1], which regulates the extent to which affective carry-over biases choice. Higher values of *φ*_*t*_ correspond to stronger regulatory engagement, thereby attenuating the influence of affective states on subsequent decisions while η_φ_ governs updating of the reappraisal state in response to the demand of contextual changes.

Following each decision, the model evaluates the selected action relative to the alternative in value space. The appraisal signal is defined as:

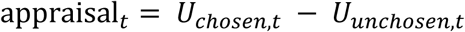

This signal is decomposed into:

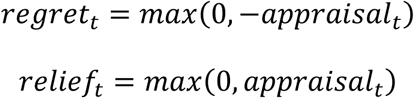

capturing adverse and favorable post-choice evaluations, respectively.

Affective intensity is computed as:

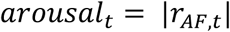

The trial-wise reappraisal demand is then given by:

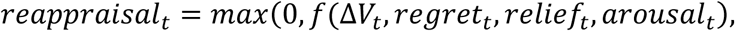

Where f(.) represents a linear function with fixed weights of each component. As a result, reappraisal_t_ increases with affective discrepancy, regret, and outcome intensity. This demand is transformed into a target reappraisal state:

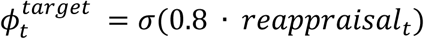

and the latent reappraisal state is updated via:

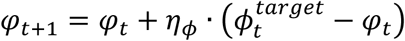

where η_ϕ_ controls the rate of adjustment.

### Contextual biases and meta-control

Beyond value-based learning, MACA-Q incorporates meta-control processes that regulate the influence of contextual and heuristic signals favoring escape. At the slower contextual level, the escape option received additive bias terms reflecting threat context and block-level escape advantage. Hence, contextual biases are defined as:

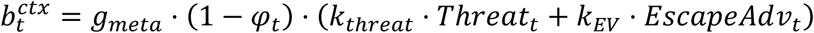

In addition to slower contextual biases, the model maintained dynamic reliance on two escape-favoring within-context heuristics: (1) an ordinal cue reflecting relative escape attractiveness (dStars), and (2) a frontbus cue signaling escape risk structure. These cues did not replace learned values; rather, they were weighted by latent reliance states, *w_stars_* (fixed parameter) and *k_risk_*. Hence, heuristic biases are defined as:

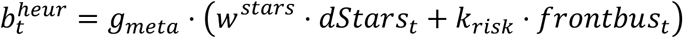

These biases acted outside the value-comparison process and therefore allowed context to influence choice independently of learned action values. Meta-control, implemented via g_meta_, scales reliance on these signals, while reappraisal *φ*_*t*_ attenuates their influence.

Meta-control and post-choice regulatory adaptation operate at distinct computational levels. Meta-control, implemented via g_meta_, scales the influence of contextual and heuristic bias signals on the decision variable at each trial, thereby determining the extent to which non-value information contributes to action selection. In contrast, the parameter η_ϕ_ governs the rate of updating of the post-choice regulatory states φ_t_. Thus, g_meta_ controls the *instantaneous expression* of contextual biases in choice, whereas η_ϕ_ controls the *dynamics of regulatory adaptation* that determine how experiences are integrated over time. These processes are complementary: meta-control sets how strongly context influences decisions, while regulatory updating determines how internal states adjust to changing task demands and outcomes.

### Parameters estimated in the winning model

The best-fitting model variant estimated the following free parameters: a base learning rate for value and success updating (*α*); baseline inverse temperature (*β*); sensitivity to structural efficacy (*k_eff_*); integration rates for latent competence (*η_SE_*), affective updating (*η_AF_*), and reappraisal/endorsement updating (*η_φ_*); a direct threat bias on escape choice (*k_threat_*); baseline regulatory gain (*Φ*); a block-level escape-advantage bias (*k_EV_*); a positive meta-control gain scaling contextual and heuristic biases (*g_meta_*); and a frontbus cue-bias parameter (*k_frontbus_*). Parameters were estimated on unconstrained scales in Stan and transformed to their appropriate bounded domains before model evaluation. The winning specification additionally included all major dynamic modules: threat-sensitive affective updating, affective carry-over into choice, effort costs, expected-success learning, agency-gated persistence learning, dynamic gain modulation by latent competence, contextual bias gating by reappraisal, dynamic reappraisal and endorsement, constructive and inertial sunk-cost routing, and surprise-gated updating of heuristic reliance. We provide a full list of parameters and latent state variables in Supplementary Table S14 and provide a comprehensive model comparison in Supplementary Table S16.

### Model Summary

MACA-Q decomposes persistence–escape decisions into separable but interacting latent processes. Standard reinforcement learning captures evolving action values for persistence and escape. A reference-dependent affective system tracks whether outcomes feel better or worse than recent experience and stores separate emotional carry-over for persistence and escape. A competence system tracks expected success, trial-wise structural efficacy, and a slower latent self-evaluative state that sharpens value-based choice when persistence is repeatedly effective. An agency module determines whether persistence outcomes should update value, thereby distinguishing failures that are informative about controllable performance from failures that appear decoupled from effort. Finally, a meta-control system dynamically regulates how strongly contextual and heuristic cues bias escape, with bias expression increasing under surprise and contextual transitions but decreasing when reappraisal engagement is high.

### Computational Fitting Procedure

Parameters of MACA-Q were estimated in Stan using R from trial-by-trial choice data.^105^ Because MACA-Q includes multiple interacting latent processes with trial-wise state updates, we used a computationally tractable approximate hierarchical procedure rather than fitting a fully joint multilevel model. Specifically, we adopted a two-stage empirical-Bayes (known as “poor man’s hierarchical”) approach that is widely used in reinforcement learning domains and computational psychiatry to obtain population-informed shrinkage when full hierarchical estimation is computationally expensive.^108–111^ In this approach, individual parameters are first estimated separately under weakly informative priors, and the resulting sample distribution is then used to construct empirical priors that regularize a second round of subject-level estimation. Such hierarchical shrinkage has been shown to improve point estimates and reduce overfitting relative to fully independent estimation, while remaining more tractable than full joint sampling for complex models.^112^

Concretely, model fitting proceeded in two rounds. In Round 1, each participant was fit separately using maximum-a-posteriori optimization with weakly informative priors in an unconstrained raw parameter space. Parameters were optimized in Stan with gradient-based L-BFGS. To reduce sensitivity to local failures, each fit was attempted from multiple initializations, and unsuccessful optimizations were re-run with a Newton fallback. Participant-level raw-parameter estimates from usable Round-1 fits were then aggregated to construct a group-level multivariate normal empirical prior, defined by the sample mean vector and covariance matrix in raw space. The covariance matrix was regularized by ridge adjustment to ensure positive definiteness, and its scale was mildly inflated to avoid overly aggressive shrinkage. In Round 2, each participant was refit under this population-informed prior. This yields regularized MAP estimates that borrow strength from the group while preserving individual variation. Final interpretable parameters were obtained by deterministic transformation of the optimized raw parameters to their constrained ranges.

We emphasize that this is an established procedure for approximating full hierarchical Bayesian inference.^108–111^ Moreover, we did not rely on fitted latent variables alone for interpretation. Instead, the adequacy of the model and the interpretability of its latent processes were evaluated using predictive checks, systematic ablation analyses, and semi-empirical parameter-recovery analyses reported in the Supplement and explained next.

### Computational Model Validation and Robustness Checks

To evaluate the adequacy and practical identifiability of MACA-Q, we conducted three complementary validation and robustness analyses:

### Predictive checks

To assess whether the MACA-Q model can account for the observed empirical data, we performed simulation-based predictive checks shown in Fig 4A-E and in the Supplement. These analyses evaluate the model’s generative validity by testing whether fitted parameter estimates reproduce the structured behavioral patterns observed in the data beyond individual-level goodness-of-fit. For each participant, we replayed their observed task sequence using their fitted parameter estimates obtained from the empirical Bayes (MAP) procedure and computed trial-wise predicted choice probabilities. This procedure preserves the full context-dependent structure of the paradigm, including incentive regimes, escape contingencies, and effort manipulations. These model-derived predictions were compared to the empirical data at both the trial-by-trial level and in terms of condition-wise choice structure. This allowed us to evaluate whether the model captures the principal qualitative and quantitative signatures of behavior. To quantify uncertainty in aggregate patterns, we used subject-level bootstrap resampling. As a robustness check, we additionally performed forward simulations in which behavior was generated from the fitted model without conditioning on the observed trial sequence. These yielded qualitatively similar results. Additional robustness analyses demonstrating that the model reproduces the observed behavioral structure are reported in the Supplement.

### MACA-Q module ablation analysis

We conducted a systematic ablation study across a broad set of nested and component-altered model variants (see Table S16). This included more than 20 theoretically motivated reduced models derived from MACA-Q. Each variant was fit after removing, fixing, or selectively disabling specific computational components to assess their contribution to model performance. This allowed us to assess which model components materially contributed to explanatory performance, whether key behavioral signatures depended on the proposed latent mechanisms, and whether improvements in fit reflected structured computational assumptions rather than diffuse model flexibility. To this end, pseudo-subject parameters were drawn from empirical fit distributions, behavior was simulated under each model variant, and models were refit using the same hierarchical estimation pipeline applied to the empirical data. This design enabled evaluation of both predictive performance and parameter recoverability under realistic parameter regimes, providing a controlled test of the functional contribution of individual components. Ablations were implemented as functional suppressions within a shared parameterization rather than as full structural removals. This choice reflects the fact that key computational processes in MACA-Q are inherently intertwined within a hierarchical control architecture, such that removing individual components would require broader reparameterization and alter the interpretation of remaining parameters. Consistent with this, we show that simpler models lacking these integrated mechanisms, such as standard Q-learning, fail to capture core behavioral signatures of the task. Within this context, maintaining a common parameterization allows us to isolate the contribution of individual components while minimizing confounds arising from differences in optimization behavior. Additional methodological details and full results of these analyses are provided in the Supplement.

### Parameter recovery analysis

We conducted semi-empirical parameter recovery analyses to assess practical identifiability and separability of model parameters. Synthetic datasets were generated by sampling parameter vectors from the empirical joint posterior distribution of fitted parameters (N = 400 simulated agents per model). For each model, subject-level parameter estimates from the empirical data were used to approximate the multivariate parameter distribution from which new parameter sets were drawn using a bootstrap-based sampling procedure. For each sampled parameter vector, behavioral data were simulated using the same task structure as in the empirical study. The resulting synthetic datasets were then refit using the identical estimation pipeline applied to the empirical data, including the same optimization procedures and parameter transformations. Parameter recovery was evaluated by comparing generating (“true”) and recovered parameter values at the subject levels. Recovery performance was quantified using Pearson correlations, Spearman correlations, root mean squared error (RMSE), and bias (mean and standard deviation of estimation error). Analyses were conducted in both raw parameter space and constrained space to ensure robustness across parameterizations.

Across primary model parameters, recovered estimates closely tracked generating values, indicating robust identifiability. In contrast, recovery was reduced for a subset of auxiliary parameters, which primarily capture slower meta-learning dynamics (η parameters). These parameters require greater temporal depth to be reliably estimated and are therefore more sensitive to data limitations. Importantly, model comparison and ablation analyses provided in the Supplement demonstrated that excluding these parameters impaired model fit. This supports their inclusion as part of the full generative architecture despite reduced recoverability at the individual parameter level. Together, these analyses confirm that the model can reliably recover the core computational mechanisms of interest while appropriately capturing additional modulatory processes that operate at longer timescales or under specific contextual conditions.

### MACA-Q Analyses of Computational Pathways into Avoidance

To characterize the computational pathways into avoidance, we conducted targeted simulation analyses using the MACA-Q model. Specifically, key model parameters were systematically varied while holding all other parameters fixed. Key parameters that we varied included: threat sensitivity (*k_threat_*), reappraisal attunement (η_φ_), meta-control strength (gₘₑₜₐ), and feedback responsiveness (α), and other parameters reported in the Supplement. For each parameter, synthetic cohorts were generated by sampling values across a predefined range (i.e., +/- 1SD from the mean of the observed empirical distribution) and simulating behavior under the full task structure. Resulting behavioral and latent trajectories were summarized at the subject level to quantify cross-context relationships between agency dynamics and escape behavior. For visualization in Fig 5, simulated subjects were stratified into high versus low parameter regimes based on median splits. These simulation-based analyses isolate the causal contribution of individual computational mechanisms to observed behavioral patterns. Additional details and extended analyses are provided in the Supplement.

To quantify cross-context relationships between agency dynamics and escape behavior (Fig 5D-F), we computed three subject-level summary indices from simulated data, namely: *End-of-safety agency*, which refers to the mean inferred agency over the final 5 trials of safe blocks (pS and eS), capturing accumulated agency prior to threat exposure. *Escape probability under threat* (pT): mean escape probability within threat blocks where persistence was advantageous. *Threat buffering index*: difference in escape probability between controllable and degraded-controllability threat conditions (pT_cl_ - pT), indexing adaptive sensitivity to loss of control. In addition, full trial-wise trajectories of latent variables (e.g., agency, value signals) were retained to characterize dynamic adaptation across contexts.

## Acknowledgment

We thank Michael J. Frank for insightful discussions on the results and the computational modeling analyses as well as for thoughtful feedback on intermediate stages of this project. We also thank Martin Paulus for insightful discussions and valuable feedback on earlier drafts of this manuscript. We also thank the entire Gearshift Fellowship development team, in particular Madeline Healey, Rachel Zhu, Rhuan Martinez, Rob Gemma, and Josh Lu, for implementing the paradigm.

