## Supplemental Material for "Sustaining Control and Agency Under Threat: Computational Pathways to Persistence and Escape"

### 1. Simulation details of parameter identifiability analysis

To further assess parameter identifiability, we conducted additional simulation analyses examining the effect of data quantity on recovery performance. Specifically, we varied the number of trials per block and overall task length while keeping the generative parameter distributions fixed. These analyses revealed that recovery of a subset of auxiliary parameters improved systematically with increasing trial numbers. This pattern is consistent with the functional role of these parameters: k parameters capture transient, context-dependent biases induced by heuristic cues, whereas η parameters govern slower meta-learning and meta-control dynamics. As such, their reliable estimation requires sufficient temporal depth to disentangle short-term fluctuations from longer-term adaptive processes. Importantly, these findings indicate that reduced recovery for these parameters in the present dataset reflects data limitations rather than model misspecification.

### 2. Ablation details of MACA-Q model components analysis

To evaluate the functional contribution of individual model components, we conducted a systematic ablation analysis within the MACA-Q modeling framework. The goal of this analysis was to determine which components are necessary to reproduce key behavioral signatures and to assess their impact on both predictive performance and parameter recoverability under realistic parameter regimes. Model variants were defined through a structured configuration file that specified the inclusion or suppression of individual components (see Table S16 for a summary of model comparisons). For each variant, the overall model architecture and parameterization were held constant, while targeted components were either fully active or functionally suppressed via parameter constraints or tight priors centered on neutral values. This approach allowed a controlled comparison across model variants while maintaining a unified parameter space and estimation framework. This analysis revealed that specific components, particularly those introduced in the main text and governing meta-control mechanisms, affective learning and agency inference, were critical for capturing key behavioral signatures and maintaining parameter identifiability across contexts (see Table S16 & Figures S3-S4).

Importantly, ablations were implemented as functional suppressions within a shared parameterization rather than as full structural removals of model components. Targeted modules were constrained toward neutral or non-influential regimes via parameter settings or priors, while retaining a consistent overall model architecture across variants. This approach preserves a stable estimation framework and avoids confounds arising from reparameterization or differences in optimization behavior that can accompany structurally distinct models. It is particularly appropriate in the present context, where key computational mechanisms are inherently intertwined within a hierarchical control architecture, such that removing individual components would require broader changes to the model structure and could alter the interpretation of other parameters. Consistent with this, we show in the main text that simpler models lacking these integrated mechanisms, such as standard Q-learning, fail to capture core behavioral signatures of the task. Consequently, the ablation analysis provides a controlled test of the functional contribution of individual components within a unified computational architecture. Differences between variants therefore reflect the extent to which each component contributes to predictive performance, behavioral pattern formation, and parameter identifiability, rather than differences in model flexibility arising from changes in parameterization.

To ensure that simulations were grounded in empirically plausible parameter regimes, we adopted a semi-empirical parameter generation procedure. For each model variant, empirical parameter estimates were first obtained from fits to real participant data. Pseudo-subject parameter sets were then generated by sampling from these empirical joint distributions (using bootstrap resampling). This procedure preserves both the marginal distributions and covariance structure of parameters. This ensures that simulated behavior reflects realistic variability observed in the empirical data rather than relying on arbitrary or hand-specified parameter values. For each model variant, we generated pseudo-subjects (N=400), each characterized by a full parameter vector drawn from the corresponding empirical distribution.

For each pseudo-subject, behavioral data were simulated using the same paradigm setup like in the actual empirical study and the corresponding model variant. Specifically, the simulated paradigm consisted of five blocks with twelve trials per block, yielding a total of 60 trials per subject. We used the same task structure as in the online study administered to actual subjects. Hence, the simulations preserved all key structural features of the task, including trial-by-trial variation in reward magnitude and effort, stochastic outcome generation, and context-dependent manipulations such as incentive context (safe versus threat) and changes in controllability. Across pseudo-subjects, this resulted in 24,000 simulated trials per model variant. This design ensured that simulated datasets were directly comparable across model variants and matched the scale and structure of the empirical data.

All simulated datasets were refit using the same estimation pipeline applied to the empirical data. Parameter estimation followed a hierarchical empirical-Bayes procedure. In a first stage, subject-level parameters were estimated independently using gradient-based optimization. These estimates were then used to derive group-level priors, which were subsequently incorporated into a second round of subject-level fitting to stabilize parameter estimates. Optimization was performed using L-BFGS with fallback strategies where necessary, and only valid fits passing convergence and quality checks were retained for subsequent analyses. This procedure ensured that both recovery and model comparison reflected the same estimation regime used in the primary analyses (see description in the Methods of the main manuscript).

Model variants were evaluated along two complementary dimensions: predictive performance and parameter recovery. Predictive performance was assessed using trial-level likelihood-based and calibration-based metrics computed on prediction datasets, including summed log-likelihood, Brier score, and calibration error quantified as the root mean squared error between predicted and observed choice probabilities. These metrics capture both the accuracy and calibration of model predictions and allow comparison across variants in terms of their ability to reproduce observed behavioral patterns.

Parameter recoverability was assessed by comparing true (generating) and recovered parameter values at the subject level. For each parameter, we computed Pearson correlations between true and recovered values, as well as root mean squared error (RMSE) and bias, defined as the mean and standard deviation of the estimation error (recovered minus true). Additional diagnostics included examination of error distributions, assessment of heteroscedasticity by relating absolute error to parameter magnitude, and analysis of correlation structure among recovered parameters to detect potential dependencies or trade-offs between parameters. To ensure comparability across model variants, recovery analyses were optionally restricted to parameters that were functionally active in each configuration, based on the model specification flags.

Because ablated components were suppressed within a shared parameterization rather than fully removed, the ablation analysis should be interpreted as a test of functional contribution rather than a strict nested-model comparison. Differences between model variants therefore reflect the extent to which a given component improves predictive performance, stabilizes parameter estimation, and enables recovery of interpretable latent variables under realistic parameter regimes. This distinction is particularly important given the use of hierarchical empirical-Bayes estimation, which introduces shrinkage and may influence parameter identifiability.

Together, this semi-empirical ablation framework provides a principled approach to evaluating model components by combining realistic parameter sampling, full task simulation, consistent hierarchical estimation, and multi-metric evaluation of prediction and recovery. This design enables identification of components that are not only theoretically motivated but also empirically necessary to account for observed behavior and to yield stable, interpretable parameter estimates within the MACA-Q model.

### 3. Results from Ablation analyses

We conducted a systematic ablation analysis in which key components of the MACA-Q architecture were selectively disabled and evaluated using posterior predictive checks (Fig S2). While reduced models were often able to approximate coarse block-level escape rates, clear and systematic deviations emerged in the temporal dynamics and context-dependent adaptation profiles.

Specifically, distinct ablations produced qualitatively different failure signatures. Removal of contextual differentiation mechanisms led to impaired sensitivity to task structure, particularly at the onset of escape-favorable contexts (eT, eS), where models failed to appropriately increase escape behavior. In contrast, disabling policy calibration mechanisms resulted in systematic misallocation of behavior, characterized by overprediction of escape when persistence was advantageous (pS, pT) and underprediction when escape was optimal (eT, eS). Ablations targeting global scaling processes introduced an overall upward bias in escape rates, reducing the model’s ability to reproduce context-specific differentiation.

Furthermore, removal of heuristic integration mechanisms selectively impaired sensitivity to cue-driven fluctuations, particularly in eS, where models failed to capture the transient dynamics following contextual changes. Critically, ablations affecting boundary-sensitive and control-updating processes disrupted adaptation to unannounced changes in controllability, leading to systematic overprediction of escape in the pT(cl) condition and reduced responsiveness to context transitions. Finally, removing arbitration mechanisms led to a flattening of heuristic cue differentiation within contexts, further impairing fine-grained behavioral modulation.

Together, these results demonstrate that distinct components of the MACA-Q architecture make non-redundant contributions to behavior, with each mechanism supporting specific aspects of contextual sensitivity, learning dynamics, and adaptive control. The observed dissociation of failure modes indicates that the full pattern of behavior cannot be explained by simpler or partially reduced models, but instead arises from the coordinated interaction of multiple computational processes.

Although some ablations produce superficially similar biases (e.g., increased escape in pS), they differ in their effects on within-block dynamics and context transitions, indicating distinct underlying mechanisms. Importantly, no reduced model simultaneously captured context-dependent levels, within-block adaptation, and responses to controllability shifts. Although several reduced models approximated coarse block-level averages, all showed increased predictive error relative to the full model, as reflected in higher calibration RMSE and Brier score (Table S16).

### 4. Simulation-based analyses of computational escape pathways

To characterize the computational pathways underlying escape-persistence behavior, we conducted targeted simulation analyses in which individual model parameters were systematically perturbed while holding all other parameters fixed at their baseline values.

For each parameter of interest (meta-control strength *gₘₑₜₐ*, learning rate *α*, effort sensitivity *k_eff*, and meta-learning parameters governing state updating and reappraisal, e.g., *η* terms), we generated synthetic cohorts by sampling parameter values from predefined ranges using uniform distributions. For each sampled parameter value, a synthetic subject was instantiated with all remaining parameters fixed to baseline values. For each parameter manipulation, we simulated cohorts of *N = 300* synthetic subjects. Each subject completed the full task sequence identical to the empirical paradigm, including all block structures, incentive regimes (safety vs threat), escape contingencies, and effort manipulations. Behavioral choices and latent variables were generated using the full generative model, including stochastic outcome realizations and sequential belief updating. Simulations were performed using the same task structure as in the empirical experiment, preserving: the full sequence of task blocks, trial-wise reward and risk schedules, probabilistic outcome contingencies, effort manipulations derived from stimulus difficulty. For each simulated subject, the model generated trial-by-trial choices (escape vs persistence), outcome realizations, and latent trajectories including value estimates, inferred agency, and competency signals.

For visualization purposes, simulated subjects were stratified into high and low parameter regimes using median splits on the sampled parameter values. This allowed for direct comparison of behavioral and latent trajectories across parameter regimes while preserving the underlying continuous parameter manipulation. Group-level summaries (means ± standard errors) were computed across simulated subjects within each parameter regime and condition. Relationships between agency at the end of safety contexts and subsequent escape behavior were quantified using Pearson correlations across simulated subjects. These simulation-based analyses isolate the causal contribution of individual computational mechanisms to observed behavioral patterns. By varying parameters in isolation, this approach allows identification of distinct pathways through which model components shape: the accumulation of agency, adaptation to contextual changes, and the emergence of adaptive versus maladaptive escape behavior. Importantly, because all other parameters and task structure are held constant, differences in simulated behavior can be directly attributed to the manipulated computational mechanism. Additional analyses with jittered non-focal parameters yielded qualitatively similar results, confirming that the identified pathways are robust to moderate variability in the broader parameter space.

### 5. Supplementary Figures

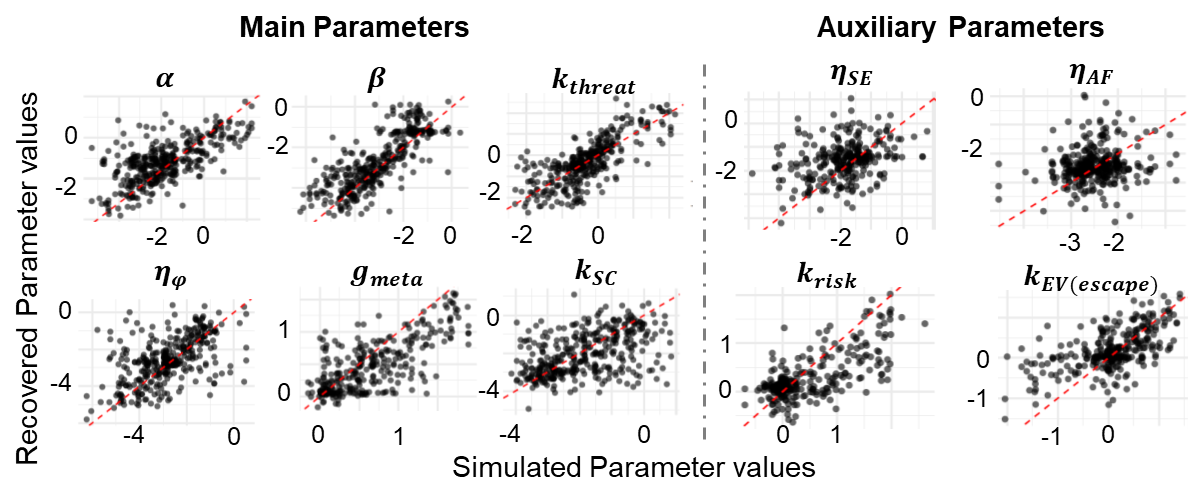

Fig. S1. Extended parameter recovery for best-fitting MACA-Q model.

Parameter recovery analyses were conducted by sampling parameters from the empirical joint posterior distribution (N=400), simulating behavioral data, and re-estimating parameters using the same fitting procedure as applied to the empirical data. Each panel shows the relationship between true (simulated) and recovered parameter values. The red dashed line denotes the identity line (perfect recovery). Across main parameters, recovered estimates closely tracked ground truth (all Pearson’s r > 0.47), indicating robust parameter identifiability. Summary statistics are reported in Table S15; parameter definitions are provided in Table S14. Parameters η correspond to meta-control and meta-learning processes governing dynamic arbitration and updating. Parameters k capture biases induced by task-related heuristic cues, including structural controllability (sc), escape risk (risk), and relative expected value of the escape option (EV_escape_; operationalized via star differences). Recovery was lower for a subset of auxiliary parameters. These parameters were not included in the main interpretation and primarily capture secondary modulatory processes. Importantly, model comparison and systematic ablation analyses showed that excluding them led to a deterioration in model fit, supporting their inclusion as part of the full generative architecture, even when individual parameter estimates are less reliably recovered. These parameters are meaningful, but require sufficient temporal depth to be resolved. Specifically, additional simulation analyses (not shown) indicate that recovery of these auxiliary parameters improves with increased trial numbers, consistent with their role in capturing transient biases (k parameters) and slower meta-learning dynamics (η parameters). This pattern suggests that their identifiability is primarily constrained by available data rather than model misspecification.

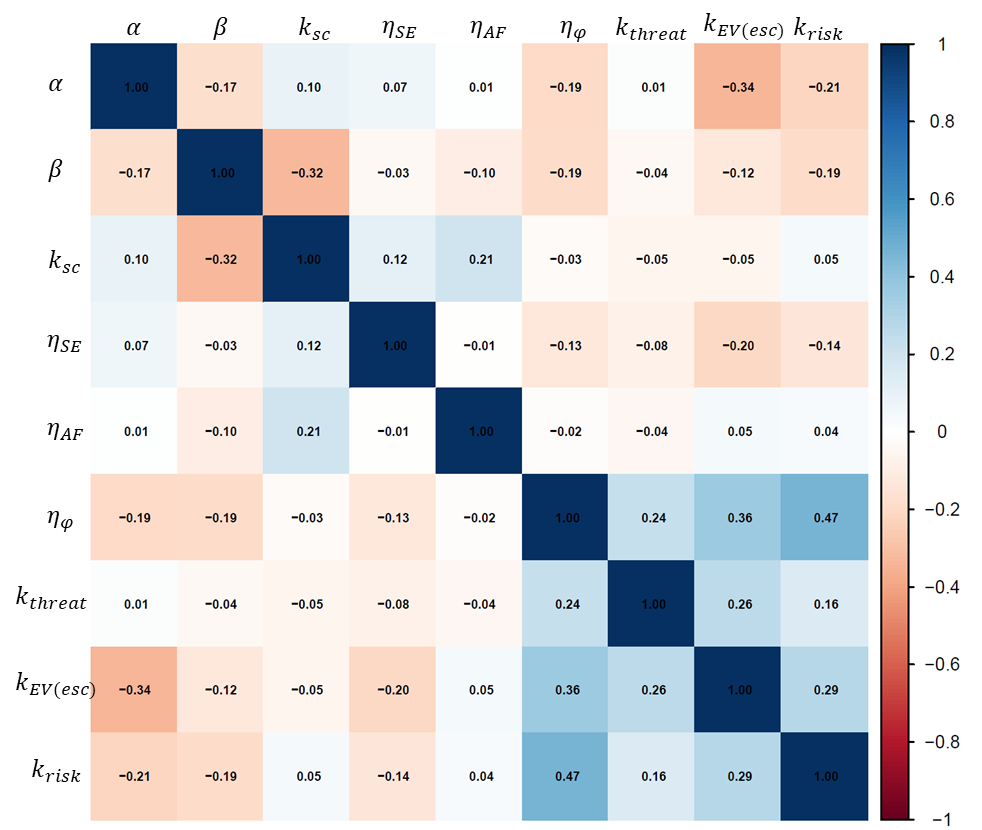

Fig. S2. Pairwise correlations between recovered MACA-Q model parameters

Overall, correlations were low to moderate, indicating partial dissociability of model components. Notably, moderate negative correlations were observed among the k parameters governing context-sensitive cue processing and the reappraisal responsiveness (η_φ_). This pattern is expected given the model architecture, in which η_φ_ dynamically modulates the influence of heuristic cues on valuation and decision-making. Thus, these dependencies reflect structured functional interactions rather than parameter redundancy. In addition, modest trade-offs were observed between learning rate (α) and cue-weighting parameters, as well as between inverse temperature (β) and several model components, reflecting well-known compensatory relationships between learning dynamics, structured valuation, and decision stochasticity in reinforcement learning models. These dependencies reflect structured functional interactions rather than parameter redundancy and did not impair the model’s ability to capture dissociable behavioral signatures, as demonstrated in the results from recovery, ablation and simulation analyses.

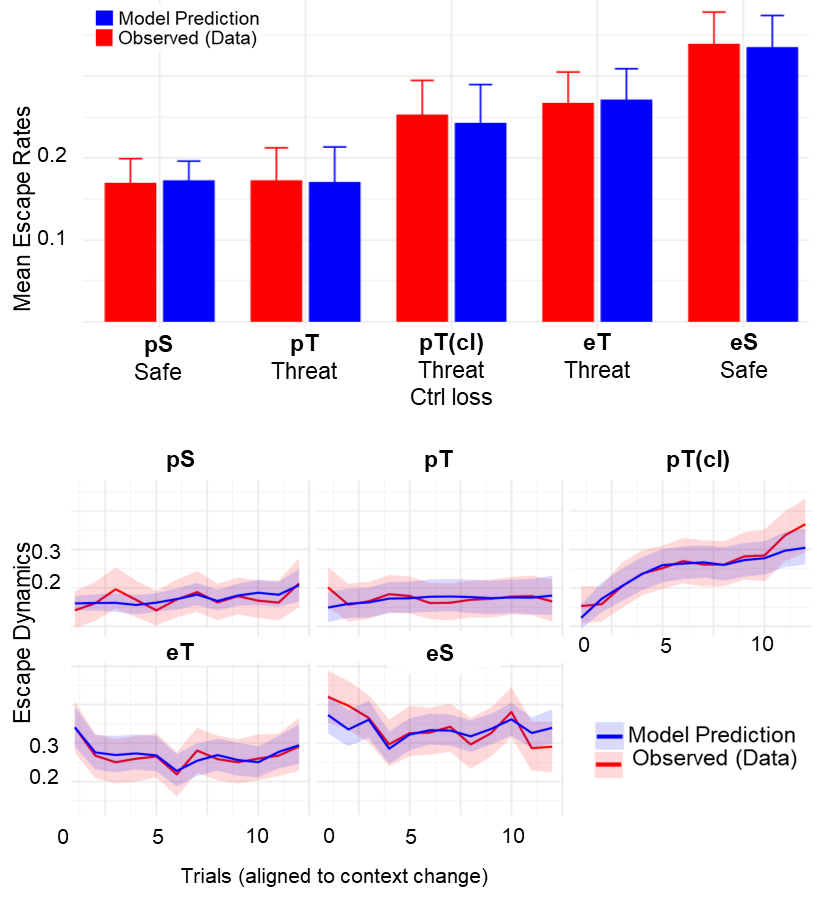

Fig. S3. Predictive checks of the MACA-Q model at both aggregate and temporal levels.

**(Top)** Model-predicted (blue) and observed (red) behavioral summaries across task conditions, showing close agreement in overall response levels. **(Bottom)** Trial-wise predictive checks across contexts. The first letter (e/p) indicates whether escape or persistence had the higher objective value (e: V(e) > V(p); p: V(e) < V(p)). The second letter (S/T) denotes the incentive regime, distinguishing safe contexts (gain vs. no gain) from threat contexts (gain vs. punishment). The pT_cl_ condition refers to a threat block in which persistence is objectively advantageous but controllability over outcomes is removed. For visualization, block contexts are shown in a fixed order. Model predictions (blue) closely track observed behavioral trajectories (red) across all conditions, capturing both the direction and shape of context-dependent dynamics over trials. This includes increasing, decreasing, and stable patterns, indicating that the model successfully reproduces structured adaptation rather than only average behavior. Minor deviations are observed primarily at the beginning of the task (e.g., early trials in eT) and toward the end of the task (e.g., late trials in pT_cl_), suggesting boundary effects related to initial adaptation and end-of-block dynamics. The model also exhibits slight smoothing of sharp transitions and modest underestimation of variability, which is expected for reinforcement learning models that integrate information over time. Overall, these results indicate that, despite its complexity, MACA-Q provides a good account of both the qualitative and quantitative structure of behavior across contexts.

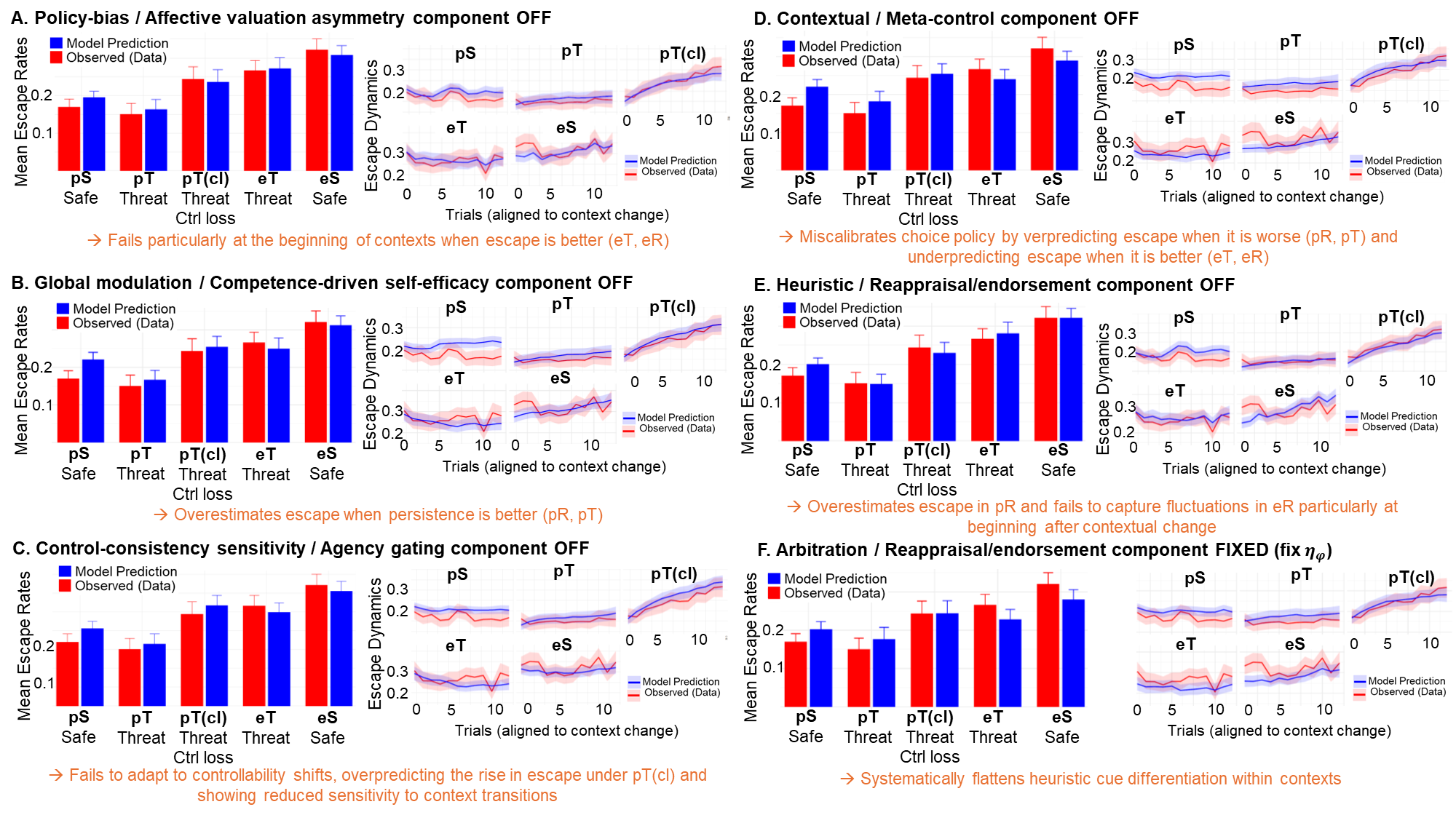

Fig. S4. Ablation analyses reveal distinct failure modes across computational mechanisms.

Shown are predictive checks (predictions in blue, empirical data in red) for representative ablated model variants. Blue indicates model predictions; red indicates observed behavior. Left panels show average escape rates across block types; right panels show trial-wise adaptation within blocks. While most ablated models retain the ability to approximate coarse block-level behavior, systematic deviations emerge in the temporal dynamics and context-dependent adaptation profiles. Several variants exhibit reduced differentiation across task contexts, overestimating escape in persistence-favorable conditions and compressing differences between pS, pT, eT, and eS blocks. Other ablations fail to capture within-block adaptation, producing overly flat trajectories that do not reflect the gradual adjustment observed in the data. Critically, multiple variants show pronounced failures in the controllability-shift condition (pT(cl)), where they fail to track the increase in escape following unannounced reductions in controllability. This pattern indicates impaired sensitivity to context transitions and disrupted integration of outcome-contingency information. Additional deviations are observed in escape conditions (eT, eS), where models lacking cue integration mechanisms fail to capture differences driven by task-relevant signals. Together, these results demonstrate that while reduced models can reproduce coarse behavioral summaries, they fail to capture distinct dynamical signatures associated with contextual adaptation, controllability inference, and cue-based modulation. These dissociable failure modes support the interpretation that multiple interacting computational mechanisms are required to account for the full structure of behavior.

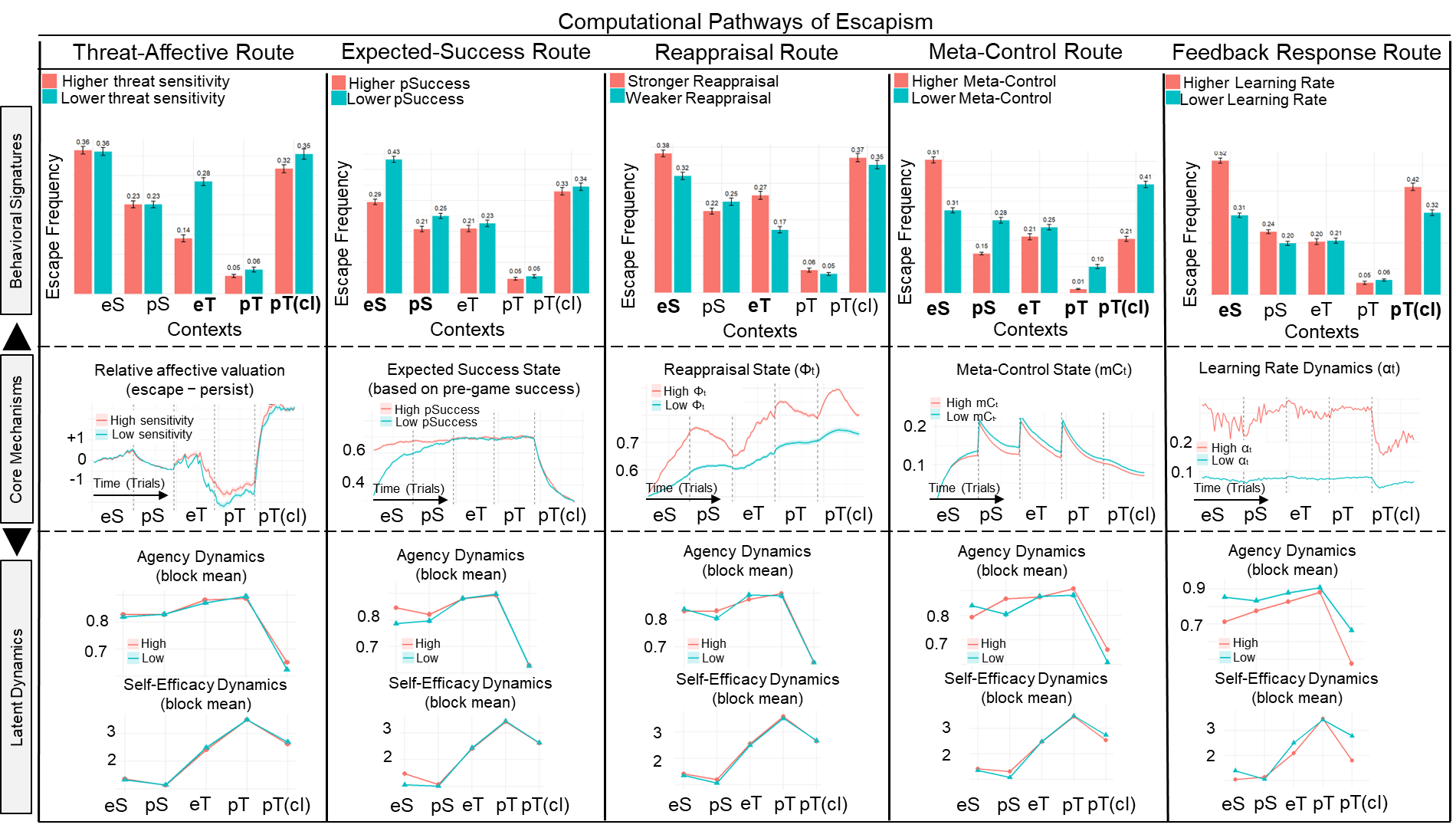

Fig. S5. Simulation-based dissociation of computational pathways with latent dynamics.

Each column shows the isolated manipulation of a single model component within its empirical range. It illustrates how distinct mechanisms give rise to similar observable behavior (top row) via different latent trajectories (middle and bottom rows). The top row displays behavioral signatures (escape probability across contexts), the middle row shows core internal signals over time, and the bottom rows depict block-wise summaries of inferred agency and self-efficacy. We refer to the Result and Method sections of the main manuscript for the approach description. Important simulation measures: 300 simulated subjects per parameter manipulation; 5 blocks of 24 trials; the sampled parameter ranges; all non-focal parameters fixed at baseline; median split for visualization. **Threat sensitivity (k_threat_) route.** Increased threat sensitivity biases outcome interpretation and policy toward persistence under threat, reducing escape despite aversive context. This effect operates at the level of valuation and policy bias without altering learning dynamics or inferred agency trajectories, consistent with a policy-level modulation of behavior. Self-efficacy increases over time when persistence remains successful, reflecting continued evidence for effective control. **Expected performance success route (p_Success_).** Higher perceived competence promotes sustained engagement and reduced escape by increasing expected success and decision precision. Self-efficacy contributes to agency formation by influencing expected competence, which is evaluated against experienced outcomes to update inferred control. When competence is high, consistent alignment between effort and outcomes supports gradual accumulation of agency. **Reappraisal responsiveness route (η_φ_).** Increased reappraisal enhances sensitivity to contextual changes, promoting adaptive switching when persistence becomes suboptimal (e.g., under controllability loss). This mechanism primarily affects outcome interpretation relative to contextual expectations and is selectively engaged at points of increased appraisal demand. Accordingly, reappraisal has only modest effects on inferred agency. **Meta-control route (g_meta_).** Higher meta-control strengthens the integration of contextual and learned signals, producing more adaptive behavior across contexts (i.e., increased escape when advantageous and reduced escape when persistence is beneficial). This mechanism stabilizes agency accumulation during reward phases by promoting consistent engagement with controllable options. Self-efficacy increases in parallel due to sustained success under stable policies. **Feedback responsiveness route** **(α).** Higher learning rates increase sensitivity to recent outcomes, leading to faster adaptation but greater volatility. This results in rapid disengagement when outcomes deteriorate and produces lower and more unstable agency estimates. Self-efficacy similarly becomes more reactive to recent feedback, reducing its stability as a signal of accumulated competence. **Self-efficacy dynamics across contexts as a relative competence signal.** In the model, self-efficacy is computed as the expected value of persistence relative to a baseline (e.g., a random or non-contingent policy). As such, it reflects the degree to which exerting control improves outcomes beyond default behavior. This formulation explains why self-efficacy can increase under threat when controllability is preserved: because baseline outcomes are worse under threat, successful persistence provides stronger evidence for effective control, amplifying perceived competence. Critically, this effect is contingent on controllability. When outcomes remain action-contingent, elevated self-efficacy supports continued engagement and contributes to agency accumulation. In contrast, when controllability is degraded, the relative advantage of persistence collapses, leading to reductions in self-efficacy and a transition toward disengagement, consistent with a pathway into helplessness. Together, these results demonstrate that persistence and escape emerge from distinct computational pathways organized through dynamically inferred agency, linking latent mechanisms to adaptive and maladaptive behavior across contexts.

### 6. Supplementary Tables

| **Characteristic** | **Value** |
| --- | --- |
| **Sample size** | N = 457 |
| **Demographics** |  |
| Age (years) | M = 31.41 (SD = 11.32), range [18-35] |
| Sex | 47% female (n = 216),  52% male (n = 240), 1%  other/prefer not to say (n = 1) |
| Ethnicity (self-identified) | White: 70% (n = 321),  Black/African American: 19% (n = 90),  American Indian/Alaska Native: 0.4% (n = 3), Asian: 6% (n = 26),  Native Hawaiian/Pacific Islander: 3% (n = 14),  Other: 0.6% (n = 4) |
| Years of education | M = 15.61 (SD = 2.92), range [6-30] |
| Highest degree | Elementary school: 0.6% (n = 2),  Middle school: 1% (n = 5),  High school: 14% (n = 65),  Some college: 21% (n = 93),  Bachelor’s: 37% (n = 169),  Master’s: 8% (n = 39),  Doctorate: 18% (n = 83),  Professional: 0.4% (n = 1) |
| **Mental health** |  |
| Mental health diagnosis  (self-reported) | 51% (n = 227) |
| **DASS-21 scores** |  |
| Depression | M = 12.90 (SD = 6.11), range [7-28] |
| Anxiety | M = 11.36 (SD = 4.57), range [7-28] |
| Stress | M = 13.31 (SD = 5.26), range [7-28] |

Table S1. Sample characteristics.

Demographic, educational, and mental health characteristics of the final sample included in the analyses. Continuous variables are reported as mean (M), standard deviation (SD), and range; categorical variables are reported as percentages with counts. Mental health diagnosis reflects self-reported lifetime diagnosis, and DASS-21 scores are presented for depression, anxiety, and stress dimensions.

| **Symptom** | **Contrast** | **Estimate** | **SE** | **Z** | **P value** | **P FDR** |
| --- | --- | --- | --- | --- | --- | --- |
| Anxiety | Framing | -0.302 | 0.097 | -3.100 | 0.0019 | 0.0023 |
| Anxiety | EV | -0.684 | 0.099 | -6.900 | 0.0000 | 0.0000 |
| Anxiety | Agency | -0.205 | 0.144 | -1.430 | 0.1530 | 0.1530 |
| Depression | Framing | 0.407 | 0.097 | 4.210 | 0.0000 | 0.0001 |
| Depression | EV | 0.540 | 0.099 | 5.480 | 0.0000 | 0.0000 |
| Depression | Agency | 0.531 | 0.144 | 3.690 | 0.0002 | 0.0003 |

Table S2. Symptom-by-contrast interaction tests with FDR correction

Estimates reflect the difference in contrast effects between high (+1 SD) and low (−1 SD) symptom levels. P values reflect nominal significance. P FDR reflect the adjusted p values using the Benjamini-Hochberg false discovery rate procedure across the six planned tests. Symptoms were measured with DASS-21 scale as detailed in the Methods of the main manuscript.

| **Dependent Variable:** | **Escape Choice** | | |
| --- | --- | --- | --- |
| *Predictors* | *Odds Ratios* | *std. Error* | *p* |
| (Intercept) | 0.164 | 0.017 | **<0.001** |
| Incentive context [threat] | 0.831 | 0.065 | **0.017** |
| Time (trial index) | 0.996 | 0.003 | 0.110 |
| **Random Effects** | | | |
| σ^2^ | 3.29 | | |
| τ_00_ _sbj_ | 3.91 | | |
| ICC | 0.54 | | |
| Observations | 21185 | | |
| Marginal R^2^ / Conditional R^2^ | 0.003 / 0.544 | | |
| AIC | 17488.001 | | |

Table S3. Mixed-effects logistic regression results assessing the robustness of the threat effect on escape behavior when controlling for time-on-task.

Escape choices were modeled as a function of incentive context (threat vs. safety) and trial index (time), with random intercepts for participants. The effect of incentive context remained significant, indicating reduced escape under threat relative to reward, even when accounting for potential time-related changes in behavior. The trial index did not significantly predict escape behavior, suggesting that the observed context effects are not explained by practice or fatigue over time. Model fit statistics (marginal and conditional R², AIC) and random effect estimates are reported for completeness. Odds ratios greater than 1 indicate increased likelihood of choosing escape.

| **Dependent Variable:** | **logRT** | | |
| --- | --- | --- | --- |
| *Predictors* | *Estimates* | *std. Error* | *p* |
| (Intercept) | 7.481 | 0.034 | **<0.001** |
| Incentive context [threat] | 0.185 | 0.032 | **<0.001** |
| Time (trial index) | -0.017 | 0.001 | **<0.001** |
| **Random Effects** | | | |
| σ^2^ | 0.39 | | |
| τ_00_ _sbj_ | 0.29 | | |
| ICC | 0.43 | | |
| Observations | 6121 | | |
| Marginal R^2^ / Conditional R^2^ | 0.071 / 0.472 | | |
| AIC | 12417.915 | | |

Table S4. Mixed-effects regression results assessing the effect of incentive framing on response times under persistence while controlling for time-on-task.

Log-transformed response times (logRT) were modeled as a function of incentive context (threat vs. safety) and trial index (time), with random intercepts for participants. Response times were significantly longer in threat contexts. Trial index showed a significant negative effect, reflecting faster responses over time, consistent with practice effects. Critically, the effect of incentive context remained robust after accounting for time-on-task, indicating that differences in response speed are not driven by temporal dynamics.

| **Dependent Variable:** | **Task accuracy under persistence** | | |
| --- | --- | --- | --- |
| *Predictors* | *Odds Ratios* | *std. Error* | *p* |
| (Intercept) | 1.512 | 0.112 | **<0.001** |
| Incentive context [threat] | 1.252 | 0.095 | **0.003** |
| Time (trial index) | 1.008 | 0.003 | **0.004** |
| **Random Effects** | | | |
| σ^2^ | 3.29 | | |
| τ_00_ _sbj_ | 1.61 | | |
| ICC | 0.33 | | |
| Observations | 16361 | | |
| Marginal R^2^ / Conditional R^2^ | 0.009 / 0.334 | | |
| AIC | 17825.049 | | |

Table S5. Mixed-effects logistic regression results assessing the effect of incentive context on task accuracy under persistence while controlling for time-on-task.

Task accuracy was modeled as a function of incentive context (threat vs. safety) and trial index, with random intercepts for participants. Accuracy was significantly higher under threat, indicating improved performance in threat contexts. Trial index also showed a significant positive effect, reflecting increasing accuracy over time, consistent with learning or practice effects. Critically, the effect of incentive context remained significant after accounting for time-on-task, indicating that enhanced performance under threat is not explained by temporal dynamics. Odds ratios greater than 1 indicate increased likelihood of accurate code cracking under persistence.

| **Dependent Variable:** | **Escape Choice** | | |
| --- | --- | --- | --- |
| *Predictors* | *Odds Ratios* | *std. Error* | *p* |
| Context [pS] | 0.072 | 0.008 | **<0.001** |
| Context [eS] | 3.540 | 0.207 | **<0.001** |
| Context [pT] | 0.845 | 0.054 | **0.009** |
| Context [eT] | 2.408 | 0.142 | **<0.001** |
| Context [pTcl] | 1.993 | 0.119 | **<0.001** |
| Anxiety z-score (ANX) | 0.928 | 0.129 | 0.590 |
| Depression z-score (DEP) | 1.362 | 0.194 | **0.030** |
| Context [eS] × ANX | 0.747 | 0.057 | **<0.001** |
| Context [pT] × ANX | 1.223 | 0.102 | **0.016** |
| Context [eT] × ANX | 0.826 | 0.063 | **0.012** |
| Context [pTcl] × ANX | 1.104 | 0.085 | 0.202 |
| Context [eS] × DEP | 1.182 | 0.088 | **0.024** |
| Context [pT] × DEP | 0.736 | 0.062 | **<0.001** |
| Context [eT] × DEP | 1.069 | 0.079 | 0.368 |
| Context [pTcl] × DEP | 0.960 | 0.073 | 0.591 |
| **Random Effects** | | | |
| σ^2^ | 3.29 | | |
| τ_00_ _sbj_ | 4.43 | | |
| ICC | 0.57 | | |
| Observations | 26120 | | |
| Marginal R^2^ / Conditional R^2^ | 0.046 / 0.593 | | |

Table S6. Mixed-effects logistic regression of escape choices as a function of task context and dimensional symptom measures.

Escape behavior was modeled as a function of task context (with pS as the reference condition), anxiety and depression (z-scored), and their interactions, with random intercepts for participants. Depression was associated with increased escape across contexts, while anxiety showed no global effect but significant context-dependent modulation. Context effects remained robust, indicating that symptom-related differences reflect altered arbitration across conditions rather than uniform shifts in behavior.

| **Dependent Variable:** | **Escape Choice** | | |
| --- | --- | --- | --- |
| *Predictors* | *Odds Ratios* | *std. Error* | *p* |
| Context [pS] | 0.071 | 0.008 | **<0.001** |
| Time (trial index) | 1.001 | 0.003 | 0.769 |
| Context [eS] | 3.536 | 0.207 | **<0.001** |
| Context [pT] | 0.827 | 0.078 | **0.045** |
| Context [eT] | 2.357 | 0.214 | **<0.001** |
| Context [pTcl] | 1.921 | 0.261 | **<0.001** |
| Anxiety z-score (ANX) | 0.925 | 0.129 | 0.576 |
| Depression z-score (DEP) | 1.364 | 0.194 | **0.029** |
| Context [eS] × ANX | 0.749 | 0.057 | **<0.001** |
| Context [pT] × ANX | 1.225 | 0.102 | **0.015** |
| Context [eT] × ANX | 0.829 | 0.063 | **0.014** |
| Context [pTcl] × ANX | 1.107 | 0.086 | 0.191 |
| Context [eS] × DEP | 1.179 | 0.087 | **0.026** |
| Context [pT] × DEP | 0.734 | 0.062 | **<0.001** |
| Context [eT] × DEP | 1.065 | 0.079 | 0.395 |
| Context [pTcl] × DEP | 0.958 | 0.072 | 0.570 |
| **Random Effects** | | | |
| σ^2^ | 3.29 | | |
| τ_00_ _sbj_ | 4.42 | | |
| ICC | 0.57 | | |
| Observations | 26120 | | |
| Marginal R^2^ / Conditional R^2^ | 0.046 / 0.593 | | |

Table S7. Mixed-effects logistic regression of escape choices including time as covariate.

Results replicated the primary model (see Table S6), with depression predicting increased escape across contexts and anxiety showing context-dependent modulation. Trial index had no significant effect, indicating that the observed patterns are not driven by time-on-task.

| **Dependent Variable:** | **Escape Choice** | | |
| --- | --- | --- | --- |
| *Predictors* | *Odds Ratios* | *std. Error* | *p* |
| Context [pS] | 0.073 | 0.008 | **<0.001** |
| Context [eS] | 0.257 | 0.028 | **<0.001** |
| Context [pT] | 0.061 | 0.007 | **<0.001** |
| Context [eT] | 0.174 | 0.019 | **<0.001** |
| Context [pTcl] | 0.144 | 0.016 | **<0.001** |
| Anxiety z-score (ANX) | 0.878 | 0.114 | 0.319 |
| Depression z-score (DEP) | 1.337 | 0.178 | **0.030** |
| Contrast [S vs T Incentive Context] × ANX | 0.918 | 0.065 | 0.224 |
| Contrast [Escape Better vs Worse Context] × ANX | 0.757 | 0.058 | **<0.001** |
| Contrast [No Control vs Control Context] × ANX | 0.786 | 0.172 | 0.271 |
| Baseline [Residual Contrast] × ANX | 1.291 | 0.153 | **0.031** |
| Contrast [S vs T Incentive Context] × DEP | 1.093 | 0.074 | 0.188 |
| Contrast [Escape Better vs Worse Context] × DEP | 1.171 | 0.087 | **0.033** |
| Contrast [No Control vs Control Context] × DEP | 1.567 | 0.338 | **0.037** |
| Baseline [Residual Contrast] × DEP | 0.784 | 0.091 | **0.036** |
| **Random Effects** | | | |
| σ^2^ | 3.29 | | |
| τ_00_ _sbj_ | 4.48 | | |
| ICC | 0.58 | | |
| Observations | 26120 | | |
| Marginal R^2^ / Conditional R^2^ | 0.045 / 0.596 | | |

Table S8. Fixed and random effects from generalized linear mixed-effects model.

Trial-wise escape decisions were modeled using a logistic mixed-effects model with a categorical predictor for contexts (encoding incentive framing, relative expected value of escape vs. persistence, and controllability), continuous symptom measures (anxiety and depression; z-scored), and their interactions. To test theory-driven hypotheses, planned contrasts corresponding to the core task dimensions [incentive framing (safety vs. threat), relative expected value (escape-favorable vs. unfavorable), and controllability (present vs. absent)] were evaluated as linear combinations of the block-level estimates within the same model. This approach allows hypothesis testing while retaining the full task structure in a unified parameterization.

| **Dependent Variable:** | **Escape Choice** | | |
| --- | --- | --- | --- |
| *Predictors* | *Odds Ratios* | *std. Error* | *p* |
| Time (trial index) | 1.000 | 0.003 | 0.907 |
| Context [pS] | 0.074 | 0.009 | **<0.001** |
| Context [eS] | 0.261 | 0.030 | **<0.001** |
| Context [pT] | 0.062 | 0.010 | **<0.001** |
| Context [eT] | 0.174 | 0.026 | **<0.001** |
| Context [pTcl] | 0.145 | 0.028 | **<0.001** |
| Anxiety z-score (ANX) | 0.879 | 0.114 | 0.317 |
| Depression z-score (DEP) | 1.334 | 0.177 | **0.030** |
| Contrast [S vs T Incentive Context] × ANX | 0.907 | 0.064 | 0.167 |
| Contrast [Escape Better vs Worse Context] × ANX | 0.742 | 0.056 | **<0.001** |
| Contrast [No Control vs Control Context] × ANX | 0.816 | 0.178 | 0.352 |
| Baseline [Residual Contrast] × ANX | 1.271 | 0.150 | **0.043** |
| Contrast [S vs T Incentive Context] × DEP | 1.097 | 0.074 | 0.170 |
| Contrast [Escape Better vs Worse Context] × DEP | 1.186 | 0.088 | **0.021** |
| Contrast [No Control vs Control Context] × DEP | 1.544 | 0.333 | **0.044** |
| Baseline [Residual Contrast] × DEP | 0.786 | 0.091 | **0.038** |
| **Random Effects** | | | |
| σ^2^ | 3.29 | | |
| τ_00_ _sbj_ | 4.41 | | |
| ICC | 0.57 | | |
| Observations | 26120 | | |
| Marginal R^2^ / Conditional R^2^ | 0.045 / 0.592 | | |

Table S9. Fixed and random effects from generalized linear mixed-effects model with time.

Same model as in Table S8 but including a continuous trial index as a covariate to account for potential practice or fatigue effects across the task. Planned contrasts of the task dimensions were evaluated as linear combinations of the block-level estimates within the same model. Results were qualitatively unchanged with the inclusion of trial index, indicating that the observed context-dependent patterns of escape behavior and their modulation by symptom dimensions cannot be explained by time-on-task effects.

| **Dependent Variable:** | **Escape Choice** | | |
| --- | --- | --- | --- |
| *Predictors* | *Odds Ratios* | *std. Error* | *p* |
| (Intercept) | 0.114 | 0.020 | **<0.001** |
| Contrast [S vs T Incentive Context] | 0.857 | 0.043 | **0.002** |
| Contrast [Escape Better vs Worse Context] | 0.762 | 0.068 | **0.002** |
| Contrast [No Control vs Control Context] | 2.964 | 0.249 | **<0.001** |
| Baseline [Residual Contrast] | 1.826 | 0.091 | **<0.001** |
| Anxiety z-score (ANX) | 0.880 | 0.118 | 0.338 |
| Depression z-score (DEP) | 1.356 | 0.186 | **0.026** |
| Effort | 1.169 | 0.074 | **0.014** |
| Reward Sensitivity | 2.086 | 0.091 | **<0.001** |
| pSuccess | 0.983 | 0.244 | 0.945 |
| Contrast [S vs T Incentive Context] × ANX | 0.909 | 0.059 | 0.139 |
| Contrast [Escape Better vs Worse Context] × ANX | 0.983 | 0.116 | 0.882 |
| Contrast [No Control vs Control Context] × ANX | 0.955 | 0.105 | 0.673 |
| Baseline [Residual Contrast] × ANX | 0.988 | 0.063 | 0.856 |
| Contrast [S vs T Incentive Context] × DEP | 1.153 | 0.073 | **0.025** |
| Contrast [Escape Better vs Worse Context] × DEP | 1.038 | 0.118 | 0.742 |
| Contrast [No Control vs Control Context] × DEP | 1.193 | 0.128 | 0.102 |
| Baseline [Residual Contrast] × DEP | 0.964 | 0.061 | 0.564 |
| ANX × effort | 1.085 | 0.085 | 0.295 |
| DEP × effort | 0.934 | 0.070 | 0.361 |
| ANX × Reward Sensitivity | 0.842 | 0.048 | **0.002** |
| DEP × Reward Sensitivity | 1.102 | 0.061 | 0.078 |
| Continue. |  |  |  |
| Continued.  **Random Effects** | | | |
| σ^2^ | 3.29 | | |
| τ_00_ _sbj_ | 4.56 | | |
| ICC | 0.58 | | |
| Observations | 26120 | | |
| Marginal R^2^ / Conditional R^2^ | 0.059 / 0.606 | | |

Table S10. Generalized linear mixed-effects model of trial-wise escape choices including within-block valuation predictors.

Trial-wise escape decisions were modeled using a logistic mixed-effects model including planned contrasts capturing the core task dimensions: incentive framing (safety vs. threat), relative expected value (escape-favorable vs. unfavorable), and controllability (present vs. absent) as well as continuous symptom measures (anxiety and depression; z-scored) and their interactions. To dissociate trial-level valuation processes from block-level contextual structure, the model additionally included effort (binary task difficulty: high vs. low), reward sensitivity (trial-wise heuristic cue derived from package features indicating potential reward magnitude), and pSuccess (participant-specific probability of successful task completion, estimated from performance in an independent paradigm; see Methods). Results indicate that both contextual factors and within-trial valuation signals contribute to escape decisions, with effort and reward sensitivity exerting significant effects, while pSuccess showed no main effect.

| **Dependent Variable:** | **Escape Choice** | | |
| --- | --- | --- | --- |
| *Predictors* | *Odds Ratios* | *std. Error* | *p* |
| (Intercept) | 0.112 | 0.022 | **<0.001** |
| Time (trial index) | 1.000 | 0.003 | 0.867 |
| Contrast [S vs T Incentive Context] | 0.872 | 0.098 | 0.223 |
| Contrast [Escape Better vs Worse Context] | 0.767 | 0.072 | **0.005** |
| Contrast [No Control vs Control Context] | 2.901 | 0.422 | **<0.001** |
| Baseline [Residual Contrast] | 1.816 | 0.108 | **<0.001** |
| Anxiety z-score (ANX) | 0.881 | 0.118 | 0.342 |
| Depression z-score (DEP) | 1.354 | 0.186 | **0.027** |
| Effort | 1.168 | 0.074 | **0.015** |
| Reward Sensitivity | 2.085 | 0.091 | **<0.001** |
| pSuccess | 0.985 | 0.245 | 0.951 |
| Contrast [S vs T Incentive Context] × ANX | 0.911 | 0.059 | 0.147 |
| Contrast [Escape Better vs Worse Context] × ANX | 0.987 | 0.116 | 0.913 |
| Contrast [No Control vs Control Context] × ANX | 0.951 | 0.105 | 0.650 |
| Baseline [Residual Contrast] × ANX | 0.987 | 0.063 | 0.832 |
| Contrast [S vs T Incentive Context] × DEP | 1.151 | 0.073 | **0.026** |
| Contrast [Escape Better vs Worse Context] × DEP | 1.034 | 0.117 | 0.768 |
| Contrast [No Control vs Control Context] × DEP | 1.196 | 0.129 | 0.097 |
| Baseline [Residual Contrast] × DEP | 0.966 | 0.061 | 0.581 |
| ANX × effort | 1.084 | 0.085 | 0.300 |
| DEP × effort | 0.934 | 0.070 | 0.363 |
| ANX × reward Sensitivity | 0.841 | 0.048 | **0.002** |
| DEP × reward Sensitivity | 1.104 | 0.061 | 0.072 |
| Continue. |  |  |  |
| Continued.  **Random Effects** | | | |
| σ^2^ | 3.29 | | |
| τ_00_ _sbj_ | 4.56 | | |
| ICC | 0.58 | | |
| Observations | 26120 | | |
| Marginal R^2^ / Conditional R^2^ | 0.059 / 0.606 | | |

Table S11. Generalized linear mixed-effects model of trial-wise escape choices including contextual contrasts, trial-level valuation predictors, and time (trial index).

Trial-wise escape decisions were modeled using a logistic mixed-effects model including planned contrasts capturing the core task dimensions together with continuous symptom measures (anxiety and depression; z-scored) and their interactions. To dissociate contextual arbitration from trial-level valuation processes, the model additionally included effort (binary task difficulty: high vs. low), reward sensitivity (trial-wise heuristic cue derived from package features indicating potential reward magnitude), and pSuccess (participant-specific probability of successful task completion estimated from performance in an independent paradigm; see Methods). A continuous trial index (time) was included to account for potential practice, fatigue, or sequential-order effects. Results remained qualitatively unchanged relative to models without time, indicating that observed effects are not driven by global temporal trends. Effort and reward sensitivity significantly influenced escape behavior, whereas pSuccess showed no main effect.

| **Dependent Variable:** | **Escape Choice** | | |
| --- | --- | --- | --- |
| *Predictors* | *Odds Ratios* | *std. Error* | *p* |
| (Intercept) | 0.100 | 0.018 | **<0.001** |
| Contrast [S vs T Incentive Context] | 0.853 | 0.043 | **0.001** |
| Contrast [Escape Better vs Worse Context] | 0.757 | 0.068 | **0.002** |
| Contrast [No Control vs Control Context] | 2.994 | 0.252 | **<0.001** |
| Baseline [Residual Contrast] | 1.836 | 0.092 | **<0.001** |
| Anxiety z-score (ANX) | 0.808 | 0.180 | 0.339 |
| Depression z-score (DEP) | 1.777 | 0.410 | **0.013** |
| Effort | 3.062 | 0.447 | **<0.001** |
| Reward Sensitivity | 2.100 | 0.092 | **<0.001** |
| pSuccess | 1.132 | 0.285 | 0.621 |
| Contrast [S vs T Incentive Context] × ANX | 0.907 | 0.059 | 0.131 |
| Contrast [Escape Better vs Worse Context] × ANX | 0.981 | 0.116 | 0.874 |
| Contrast [No Control vs Control Context] × ANX | 0.961 | 0.106 | 0.722 |
| Baseline [Residual Contrast] × ANX | 0.991 | 0.063 | 0.890 |
| Contrast [S vs T Incentive Context] × DEP | 1.156 | 0.073 | **0.022** |
| Contrast [Escape Better vs Worse Context] × DEP | 1.037 | 0.118 | 0.750 |
| Contrast [No Control vs Control Context]× DEP | 1.184 | 0.128 | 0.118 |
| Baseline [Residual Contrast] × DEP | 0.961 | 0.061 | 0.529 |
| ANX × effort | 1.869 | 0.388 | **0.003** |
| DEP × effort | 0.505 | 0.102 | **0.001** |
| ANX × reward Sensitivity | 0.845 | 0.048 | **0.003** |
| DASS Depress z × reward Sensitivity | 1.099 | 0.061 | 0.088 |
| effort1 × pSuccess | 0.185 | 0.041 | **<0.001** |
| Continue. |  |  |  |
| Continued. |  |  |  |
| ANX × effort[low] × pSuccess | 1.775 | 0.712 | 0.153 |
| ANX × effort[high] × pSuccess | 0.620 | 0.216 | 0.170 |
| DEP × effort[low] × pSuccess | 0.393 | 0.148 | **0.013** |
| DEP × effort[high] × pSuccess | 1.170 | 0.384 | 0.633 |
| **Random Effects** | | | |
| σ^2^ | 3.29 | | |
| τ_00_ _sbj_ | 4.60 | | |
| ICC | 0.58 | | |
| Observations | 26120 | | |
| Marginal R^2^ / Conditional R^2^ | 0.062 / 0.609 | | |

Table S12. Generalized linear mixed-effects model of trial-wise escape choices including contextual contrasts and higher-order interactions between effort, reward sensitivity, individual capability (pSuccess), and symptom dimensions.

Trial-wise escape decisions were modeled using a logistic mixed-effects model including planned contrasts capturing the core task dimensions together with continuous symptom measures (anxiety and depression; z-scored) and their interactions. To characterize trial-level valuation processes in greater detail, the model included effort (binary task difficulty: high vs. low), reward sensitivity (trial-wise heuristic cue derived from package features indicating potential reward magnitude), and pSuccess (participant-specific probability of successful task completion estimated from performance in an independent paradigm; see Methods), as well as higher-order interactions between these variables. In particular, interactions between effort and pSuccess were included to test whether individual capability modulates the impact of task difficulty on escape behavior, and whether these relationships are further shaped by symptom dimensions. Results indicate that effort strongly increases escape likelihood, while its interaction with pSuccess reveals differential sensitivity to difficulty depending on individual capability. Symptom-dependent interactions further suggest that anxiety and depression differentially modulate how effort and reward cues influence decision-making.

| **Dependent Variable:** | **Escape Choice** | | | |
| --- | --- | --- | --- | --- |
| *Predictors* | *Odds Ratios* | *std. Error* | | *p* |
| (Intercept) | 0.094 | 0.019 | | **<0.001** |
| Time (trial index) | 1.002 | 0.003 | | 0.527 |
| Contrast [S vs T Incentive Context] | 0.910 | 0.103 | | 0.404 |
| Contrast [Escape Better vs Worse Context] | 0.774 | 0.073 | | **0.007** |
| Contrast [No Control vs Control Context] | 2.773 | 0.405 | | **<0.001** |
| Baseline [Residual Contrast] | 1.798 | 0.107 | | **<0.001** |
| Anxiety z-score (ANX) | 0.807 | 0.179 | | 0.333 |
| Depression z-score (DEP) | 1.788 | 0.412 | | **0.012** |
| Effort | 3.069 | 0.448 | | **<0.001** |
| Reward Sensitivity | 2.098 | 0.092 | | **<0.001** |
| pSuccess | 1.128 | 0.284 | | 0.632 |
| Contrast [S vs T Incentive Context] × ANX | 0.908 | 0.059 | | 0.137 |
| Contrast [Escape Better vs Worse Context] × ANX | 0.987 | 0.116 | | 0.913 |
| Contrast [No Control vs Control Context] × ANX | 0.960 | 0.106 | | 0.712 |
| Baseline [Residual Contrast] × ANX | 0.991 | 0.063 | | 0.884 |
| Contrast [S vs T Incentive Context] × DEP | 1.154 | 0.073 | | **0.024** |
| Contrast [Escape Better vs Worse Context] × DEP | 1.031 | 0.117 | | 0.791 |
| Contrast [No Control vs Control Context] × DEP | 1.186 | 0.128 | | 0.113 |
| Baseline [Residual Contrast] × DEP | 0.962 | 0.061 | | 0.542 |
| ANX × effort | 1.902 | 0.396 | | **0.002** |
| DEP × effort | 0.495 | 0.100 | | **0.001** |
| ANX × reward Sensitivity | 0.844 | 0.048 | | **0.003** |
| DEP × reward Sensitivity | 1.101 | 0.061 | | 0.081 |
| effort × pSuccess | 0.184 | 0.041 | | **<0.001** |
| Continue. |  |  | |  |
| Continued. |  |  | |  |
| ANX × effort[low] × pSuccess | 1.802 | 0.724 | | 0.142 |
| ANX × effort[high] × pSuccess | 0.610 | 0.213 | | 0.156 |
| DEP × effort[low] × pSuccess | 0.385 | 0.145 | | **0.011** |
| DEP × effort[high] × pSuccess | 1.182 | 0.389 | | 0.611 |
| **Random Effects** | | |  |  |
| σ^2^ | 3.29 | | | |
| τ_00_ _sbj_ | 4.60 | | | |
| ICC | 0.58 | | | |
| Observations | 26120 | | | |
| Marginal R^2^ / Conditional R^2^ | 0.062 / 0.609 | | | |

Table S13. Generalized linear mixed-effects model of trial-wise escape choices including contextual contrasts, trial-level valuation predictors, time (trial index), and higher-order interactions.

Trial-wise escape decisions were modeled using a logistic mixed-effects model including planned contrasts capturing the core task dimensions together with continuous symptom measures (anxiety and depression; z-scored) and their interactions. To dissociate contextual arbitration from trial-level valuation processes, the model additionally included effort (binary task difficulty: high vs. low), reward sensitivity (trial-wise heuristic cue derived from package features indicating potential reward magnitude), and pSuccess (participant-specific probability of successful task completion estimated from performance in an independent paradigm; see Methods), along with their interactions. A continuous trial index (time) was included to account for potential practice, fatigue, or sequential-order effects. Higher-order interactions between effort and pSuccess were included to test whether the impact of task difficulty depends on individual capability, and whether these relationships are modulated by symptom dimensions. Results remained qualitatively consistent with simpler model specifications, indicating that context-dependent escape behavior and its modulation by valuation signals and symptoms are robust to the inclusion of time and higher-order interactions.

| **Parameter** | **Function** | **Description** |
| --- | --- | --- |
| $\alpha_{t}$ | Primary | **Value learning rate (dynamically modulated by inferred agency).** Controls updating of action values based on outcomes, with learning gated by the inferred degree of controllability (agency). |
| $\beta_{t}$ | Primary | **Inverse temperature (decision precision modulated by self-efficacy).** Governs the stochasticity of choices, with higher self-efficacy increasing reliance on learned values. |
| $\eta_{\varphi}$ | Primary | **Reappraisal responsiveness.** Regulates reinterpretation of contextual and affective signals, modulating both affective updating and the influence of contextual heuristic cues on decision-making. |
| $\eta_{SE}$ | Secondary | **Self-efficacy (SE) integration rate.** Controls updating of beliefs about action success probability (self-efficacy) based on recent performance. |
| $\eta_{AF}$ | Secondary | **Affective learning rate (reference-dependent).** Governs updating of affective value representations based on deviations between experienced outcomes and a dynamic affective reference point, integrating both immediate valence signals and reappraisal-mediated reinterpretation. |
| $g_{meta}$ | Primary | **Meta-control strength.** Controls arbitration between value-based learning and contextual or heuristic influences, enabling flexible adaptation across task contexts. |
| $k_{threat}$ | Primary | **Sensitivity to threat context.** Scales the impact of threat framing (loss domain) on valuation and decision policy. |
| $k_{SC}$ | Secondary | **Sensitivity to structural controllability (instrumental efficacy).** Captures sensitivity to the objective degree of action-outcome contingency in the environment. |
| $k_{risk}$ | Secondary | **Sensitivity to heuristic risk cues.** Captures reliance on external risk indicators (e.g., probability cues such as the “frontbus”). |
| $k_{EV(Escape)}$ | Secondary | **Sensitivity to heuristic reward cues.** Captures reliance on salient reward indicators (star values associated with options). |
| **Variable** | **Function** | **Description** |
| ${Agency}_{t}$ | Latent State | **Inferred agency state.** Computed from three components: (i) inferred instrumental relevance of persistence, (ii) affective deviation of outcomes relative to a dynamic reference point, and (iii) consistency between inferred action relevance and experienced feedback. Agency increases when outcomes align with expected controllability and decreases when they violate it, with each component weighted by its magnitude. |
| ${SE}_{t}$ | Latent State | **Self-efficacy belief.** Represents the dynamically updated estimate of success probability given current task demands. |
| $\varphi_{t}$ | Latent State | **Reappraisal engagement state.** Reflects the degree to which reinterpretation processes are engaged to modulate affective and contextual meaning. |
| A_Pt_, A_Et_ | Latent State | Affective states computed from the affective module. |
| Q_Pt_, Q_Et_ | Latent State | Learned Q-values for each action (persistence vs escape) |

Table S14. Description of MACA-Q model parameters and latent state variables.

Parameters are organized according to their functional role within the MACA-Q architecture, distinguishing primary mechanisms governing value learning, affective processing, and meta-control from secondary modulatory and heuristic sensitivity parameters.

| parameter | Pearson r | rmse | Mean(bias) | SD(bias) |
| --- | --- | --- | --- | --- |
| ***β*** | 0.7975 | 0.9248 | 0.3353 | 0.8630 |
| $\boldsymbol{g}_{\boldsymbol{meta}}$ | 0.7780 | 0.3579 | -0.1409 | 0.3294 |
| $\boldsymbol{k}_{\boldsymbol{threat}}$ | 0.7618 | 0.6722 | 0.0132 | 0.6730 |
| $k_{risk}$ | 0.7228 | 0.4518 | -0.1113 | 0.4385 |
| ***α*** | 0.6603 | 1.2575 | 0.3572 | 1.2073 |
| $k_{EV(Escape)}$ | 0.5892 | 0.5176 | 0.10758 | 0.5069 |
| $\boldsymbol{\eta}_{\boldsymbol{\varphi}}$ | 0.5429 | 1.1872 | 0.2223 | 1.1677 |
| $k_{SC}$ | 0.4734 | 1.2016 | -0.1733 | 1.1906 |
| $\eta_{SE}$ | 0.3383 | 0.9126 | 0.2275 | 0.8849 |
| $\eta_{AF}$ | 0.1593 | 0.6956 | 0.0674 | 4.0000 |

Table S15. Parameter recovery statistics for the best-fitting MACA-Q model.

Recovery performance is summarized using Pearson correlation (r), root mean squared error (RMSE), and bias (mean and standard deviation of estimation error). Main model parameters (highlighted in bold font) show strong recovery, while some of the auxiliary parameters show reduced identifiability. This is consistent with their role in capturing transient biases and slower meta-learning dynamics (see Fig. S1 and Supplementary Methods). Lower RMSE indicates more accurate recovery, although interpretation should be made relative to parameter scale and in conjunction with correlation-based metrics. This pattern is consistent with hierarchical models in which processes operate across multiple timescales, with higher-order parameters requiring greater temporal depth for reliable estimation.

| **Rank** | **Model** | **Ablation type** | **Focus of**  **ablation** | **Computational interpretation** | **Calib**  **rmse** | **brier** | **Mean**  **abs_resid** | **Mean**  **loglik** | **Sum**  **loglik** |
| --- | --- | --- | --- | --- | --- | --- | --- | --- | --- |
| 1 | M1 | Reference (Best-fitting) | MACA-Q benchmark | Full integrated MACA-Q architecture | 0.2694 | 0.0726 | 0.1594 | -0.2365 | -5966.41 |
| 2 | M2 | single-module | reappraisal needed after contextual change | Contextual transitions increase perceived need for control updating | 0.2695 | 0.0726 | 0.1594 | -0.2366 | -5969.31 |
| 3 | M3 | single-module | threat modulates affective regulation | Threat changes the rate of affective updating rather than only shifting policy | 0.2699 | 0.0728 | 0.1596 | -0.2368 | -5933.31 |
| 4 | M4 | family-  level | Competence-driven efficacy & agency components | Jointly removes learned competency-driven efficacy updates and agency-gated learning (α) | 0.2698 | 0.0728 | 0.1607 | -0.2372 | -6038.89 |
| 5 | M5 | single-module | Affective carryover into utility | Learned affect feeds back into utilities at choice time | 0.2703 | 0.073 | 0.1598 | -0.2377 | -5996.14 |
| 6 | M6 | single-module | Agency-gated learning | Learning from Target outcomes is gated by inferred agency rather than updated uniformly | 0.2701 | 0.0729 | 0.1608 | -0.2377 | -6009.29 |
| 7 | M7 | single-module | Inertia sunk-cost pathway | Past investment promotes perseverative carryover independent of constructive reinterpretation | 0.2703 | 0.0731 | 0.1606 | -0.2381 | -6062.42 |
| 8 | M8 | single-module | reappraisal sensitivity after contextual change | Contextual transitions boost reappraisal responsiveness to speed adaptation (η_Φ_) | 0.271 | 0.0735 | 0.1615 | -0.2386 | -5989.21 |

Continue.

| **Rank** | **Model** | **Ablation type** | **Focus of**  **ablation** | **Computational interpretation** | **Calib**  **rmse** | **brier** | **Mean**  **abs_resid** | **Mean**  **loglik** | **Sum**  **loglik** |
| --- | --- | --- | --- | --- | --- | --- | --- | --- | --- |
| 9 | M9 | family-  level | Reappraisal / endorsement module | Reappraisal-like routing transforms regret, relief, and failure into control updates | 0.2712 | 0.0735 | 0.1585 | -0.2392 | -6202.56 |
| 10 | M10 | single-module | Affect learning | Affective states are learned from outcomes and carried forward | 0.2714 | 0.0737 | 0.1615 | -0.2395 | -6080.35 |
| 11 | M11 | family-level | Affective valuation module | Jointly removes affect learning and its carryover of biases into choice | 0.2725 | 0.0743 | 0.1626 | -0.2409 | -6102.73 |
| 12 | M12 | mechanism-variant | Heuristic trigger rule | Compares surprise-gated versus conflict-gated heuristic engagement | 0.2728 | 0.0744 | 0.1633 | -0.2411 | -5178.88 |
| 13 | M13 | single-module | Dynamic beta / precision control | Latent competency-driven self-efficacy signal dynamically tunes choice precision (β) | 0.2766 | 0.0765 | 0.1667 | -0.2475 | -6143.14 |
| 14 | M14 | single-module | within-context heuristic cue: escape advantage | Contextual escape advantage acts directly as a policy bias beyond trial value | 0.2823 | 0.0797 | 0.1712 | -0.2546 | -6454.06 |
| 15 | M15 | single-module | Threat policy bias | Threat-linked policy bias beyond learned values | 0.2838 | 0.0805 | 0.1773 | -0.2629 | -6709.32 |
| 16 | M16 | single-module | within-context heuristic cue: escape risk | Frontbus cue provides an additional learned heuristic channel about escape risk | 0.2916 | 0.085 | 0.1804 | -0.268 | -6866.49 |

Continue.

| **Rank** | **Model** | **Ablation type** | **Focus of**  **ablation** | **Computational interpretation** | **Calib**  **rmse** | **brier** | **Mean**  **abs_resid** | **Mean**  **loglik** | **Sum**  **loglik** |
| --- | --- | --- | --- | --- | --- | --- | --- | --- | --- |
| 17 | M17 | single-module | Trial-wise heuristic gating | Heuristics are engaged dynamically trial-by-trial rather than only at announced contextual changes | 0.2947 | 0.0869 | 0.1859 | -0.2762 | -7112.7 |
| 18 | M18 | single-module | within-context heuristic cue: reward sensitivity | Star-difference cues provide a learned heuristic prior for escape vs persistence | 0.2953 | 0.0872 | 0.1866 | -0.2778 | -7235.34 |
| 19 | M19 | family-level | Contextual meta-control module | Jointly removes the subsystem that scales and routes contextual bias signals | 0.3053 | 0.0932 | 0.1954 | -0.2914 | -7335 |
| 20 | M20 | single-module | Meta-control gain | Meta-control globally amplifies or attenuates contextual and heuristic influences | 0.3047 | 0.0928 | 0.2002 | -0.2934 | -7525.12 |
| 21 | M21 | family-level | Heuristic cue family | Jointly removes cue-based heuristic channels and their trial-wise recruitment | 0.3083 | 0.0951 | 0.198 | -0.2972 | -7635.66 |

Table S16. Systematic ablation analysis of MACA-Q model variants.

Overview of ablated model variants relative to the full MACA-Q model (M39), including the computational component altered, its functional interpretation, and the primary inference supported by each ablation. Predictive performance is summarized using calibration root mean squared error (RMSE) and Brier score, along with their differences relative to the full model (Δ vs M39). While several reduced models approximate coarse block-level behavior, all exhibit increased predictive error relative to the full model and, critically, show distinct and dissociable deviations in context-dependent adaptation and temporal dynamics (see Fig. S4). These results indicate that no single reduced model captures the full pattern of behavior, supporting the interpretation that adaptive escape–persistence decisions arise from the coordinated interaction of multiple computational mechanisms.
